# Genetic identification Of brain cell types underlying schizophrenia

**DOI:** 10.1101/145466

**Authors:** Nathan G. Skene, Julien Bryois, Trygve E. Bakken, Gerome Breen, James J Crowley, Héléna A Gaspar, Paola Giusti-Rodriguez, Rebecca D Hodge, Jeremy A. Miller, Ana Muñoz-Manchado, Michael C O’Donovan, Michael J Owen, Antonio F Pardiñas, Jesper Ryge, James T R Walters, Sten Linnarsson, Ed S. Lein, Major Depressive Disorder Working Group of the Psychiatric Genomics Consortium, Patrick F Sullivan, Jens Hjerling-Leffler

## Abstract

With few exceptions, the marked advances in knowledge about the genetic basis for schizophrenia have not converged on findings that can be confidently used for precise experimental modeling. Applying knowledge of the cellular taxonomy of the brain from single-cell RNA-sequencing, we evaluated whether the genomic loci implicated in schizophrenia map onto specific brain cell types. The common variant genomic results consistently mapped to pyramidal cells, medium spiny neurons, and certain interneurons but far less consistently to embryonic, progenitor, or glial cells. These enrichments were due to distinct sets of genes specifically expressed in each of these cell types. Many of the diverse gene sets associated with schizophrenia (including antipsychotic targets) implicate the same brain cell types. Our results provide a parsimonious explanation: the common-variant genetic results for schizophrenia point at a limited set of neurons, and the gene sets point to the same cells. While some of the genetic risk is associated with GABAergic interneurons, this risk largely does not overlap with that from projecting cells.

---

Knowledge of the genetic basis of schizophrenia has markedly improved in the past five years ^1^. Much of the genetic basis and heritability of schizophrenia is due to common variation ^2,3^. Identifying “actionable” genes in sizable studies ^4,5^ has proven difficult with a few exceptions ^6−8^. There is aggregated statistical evidence for diverse gene sets including genes expressed in brain or neurons ^3,5,9^, genes highly intolerant of loss-of-function variation ^10^, synaptic genes ^11^, and genes whose mRNA bind to FRMP ^12^ (Table S1). Several gene sets have been implicated by fundamentally different types of genetic studies of schizophrenia, and this convergence strongly implicates these gene sets in the pathophysiology of schizophrenia. However, the implicated gene sets often contain hundreds of functionally distinctive genes that do not immediately suggest reductive targets for experimental modeling.

Connecting the genomic results to cellular studies is crucial since it would allow us to prioritize for cells fundamental to the genesis of schizophrenia. Enrichment of schizophrenia genomic findings in genes expressed in macroscopic samples of brain tissue has been reported ^3,13,14^, but these results are insufficiently specific to guide subsequent experimentation.

A more precise approach has recently become feasible. Single-cell RNA-sequencing (scRNAseq) has been increasingly used to derive empirical taxonomies of brain cell types. We thus rigorously compared genomic results for schizophrenia to brain cell types defined by scRNAseq. We assembled a superset of brain scRNAseq data from Karolinska Institutet (KI; ***Supplementary Methods)***. Briefly, these data were generated using identical methods from the same labs with the use of unique molecular identifiers that allow for direct comparison of transcription across regions. The KI mouse superset of 9,970 cells allows identification of 24 Level 1 brain cell types and 149 Level 2 cell types, far more cell types than any other brain scRNAseq or single nuclei RNA-seq (snRNAseq) dataset now available (Figure 1A). Brain regions in the KI superset include neocortex, hippocampus, hypothalamus, striatum, midbrain, plus samples enriched for oligodendrocytes, dopaminergic neurons, and cortical parvalbuminergic interneurons (Table S2 and Figure S1). The cell type identities (both level 1 and 2) were used as they were annotated in the individual data sets –briefly cell-classification is based on clustering of cell by algorithms taking all cell class-specific gene-expression into account in a hierarchical manner. Clustering thus does not rely on the use of single markers but on the correlation of hundreds of genes. Once clustered, the cell types are identified *post-hoc* using known expression patterns and/or molecular investigation (see Table S2 for details). While most human single-cell data comes from snRNAseq, an advantage of the KI superset, and using the mouse as a model system, is that it allows for the use of scRNAseq. Prepared nuclei lack the cytoplasmic compartment and proximal dendrites, and we reasoned that this might result in a specific loss of signal. Indeed, by comparing multiple snRNAseq and scRNAseq data sets (both mouse and human), we found that transcripts destined for export to synaptic neuropil ^15^, which are enriched for genetic associations with schizophrenia (*P*=1.6x10-^4^), were significantly better captured by scRNAseq and specifically depleted in the snRNAseq (Figure S2). This unfortunately suggest that nuclear data might never provide the signal necessary for these kind of analyses.

**Figure 1.**
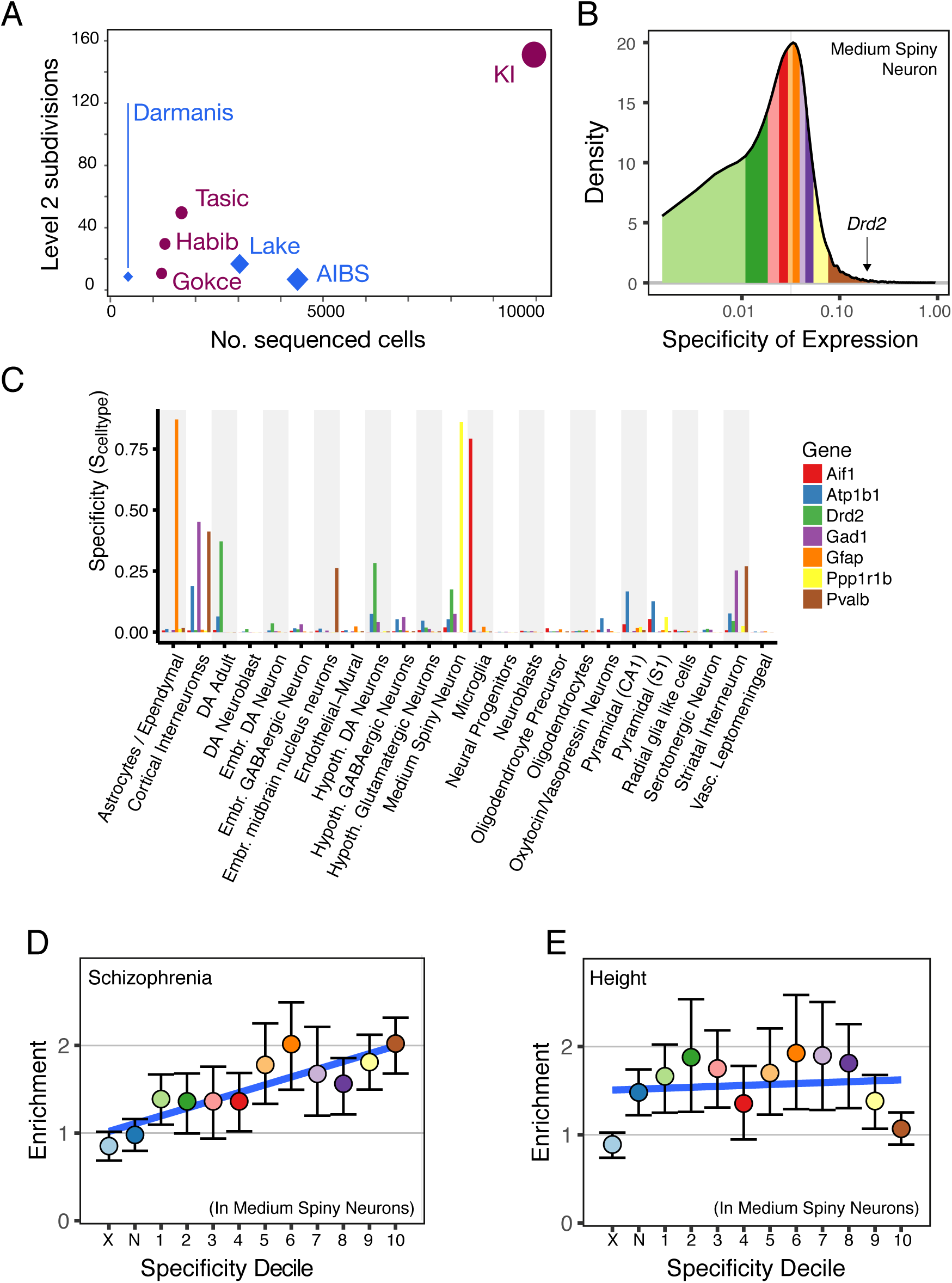
Specificity metric calculated from single cell transcriptome sequencing data can be used to test for increased burden of schizophrenia heritability in brain cell types. (A) Comparison of Level 2 cell type categories and number of cells with snRNAseq or scRNAseq from adult brain tissue (excluding retina). Circles (plum) are mouse studies and diamonds (blue) are human studies. See Table S2 for citations. AIBS=Allen Institute for Brain Science. KI=Karolinska Institutet. (B) Histogram of specificity metric (S_MSN,KI_) for medium spiny neurons from the KI superset level 1. Colored regions indicate deciles (the brown region contains the genes most specific to MSNs). Specificity value for dopamine receptor D2 *(Drd2*, S_MSN,KI,Drd2=0.17_) is indicated by the arrow. (C) Specificity values in the KI level 1 dataset for a range of known cell type markers. (D) Enrichment of schizophrenia heritability in each of the specificity deciles for medium spiny neurons (calculated using LDSC). Color of dots corresponds to regions of the specificity matrix in B. Error bars indicate the 95% confidence intervals. The light blue dot (marked ‘X’) represents all SNPs which map onto named transcripts which are not MGI annotated genes or which map onto a gene which does not have a 1:1 mouse:human homolog. The dark blue dot (marked ‘N’) represents all SNPs which map onto genes not expressed in MSNs. Blue line slows the linear regression slope fitted to the enrichment values. (E) Enrichment of height heritability in each of the specificity deciles for MSNs.

For each single-cell/nucleus data set, we estimated a specificity matrix for each gene and cell type. This measure represents the proportion of the total expression of a gene found specifically in a cell type as compared to all cell types. It is calculated as the average expression in one cell type divided by the total average expression in the other cell types. Thus, if a particular gene’s expression (even if very high) is shared between more than one cell type it will get a lower specificity measure. As an example, we show *Drd2*, a gene highly expressed in medium spiny neurons (MSNs), adult dopaminergic neurons, and hypothalamic interneurons, and thus only gets a specificity measure in MSNs of 0.17 but still ends up in the top specificity decile for that cell type (Figure 1c). To further visualize this metric, we plotted the cell type specificity values for seven genes with known expression patterns (Figure 1c). For example, the pan-neuronal marker *Atp1b1* gets lower values than specific markers *Ppp1rb1* (Darpp-32, a MSN marker), *Aif1* (Iba1, a microglia marker) and *Gfap* (an astrocyte marker) since the expression signal is spread out over several classes. For each cell type, we sorted each gene into ranked groups (i.e., deciles or 40 quantiles). The hypothesis underlying our approach is that if schizophrenia is associated with a particular cell type, then a higher amount of heritability should be located in more specific deciles/quantiles. As an example we plotted the enrichment of heritability for schizophrenia and human height in the different enrichment-deciles of MSNs, a neuronal cell type (Figure 1d,e).

To identify brain cell types associated with schizophrenia, we used the largest available genome-wide association (GWA) study of schizophrenia: CLOZUK identified ~140 genome-wide significant loci in 40,675 cases and 64,643 controls ^16^. We first compared the CLOZUK results to GTEx (RNA-seq of macroscopic samples from multiple human tissues) ^17^ and confirmed ^3^ that smaller schizophrenia GWA *P*-values were substantially enriched in brain and pituitary (Figure S3). We then evaluated the relation of the CLOZUK GWA schizophrenia results to the 24 KI Level 1 brain cell types. We applied two statistical methods, based on different assumptions and algorithms, to evaluate the association of cell type specific expression and schizophrenia common variant genetic findings. One method assessed enrichment of the common variant heritability of schizophrenia only in the most cell type-specific genes. The other method evaluated whether heritability linearly increases along with cell type expression specificity. We used two separate algorithms— LDSC^9^ and MAGMA^18^—to calculate heritability enrichments both of which accounts for confounders such as gene size and linkage disequilibrium. LDSC was used to test for enrichments in the most specific decile, while MAGMA was used to test for linear increases. We required that the two methods give similar results after correcting for multiple comparisons to minimize the chance of a spurious conclusion.

Both methods strongly highlighted hippocampal CA1 pyramidal cells, striatal medium spiny neurons, neocortical somatosensory pyramidal cells, and cortical interneurons (Figs. 2a, S4, S5). Each exceeded a stringent significance level by several orders of magnitude. The results are not pan-neuronal as multiple other types of neurons did not show enrichment, neither does the total number of molecules detected per cell type or total number of cells detected per cell type confound the results (Table S3). Schizophrenia risk was greater in mature cells than in embryonic or progenitor cells. We extended the analysis to 149 KI Level 2 cell types (Figure S6): for hippocampal CA1 pyramidal cells, both major subgroups were significant; for striatum, medium spiny neurons expressing *Drd2, Drd1* and striatal *Pvalb*-expressing interneurons were consistently significant; and for neocortical somatosensory pyramidal cells, cortical layers 2/3, 4, 5, and 6 were significant. The cortical Level 1 interneuron signal appeared to result from four Level 2 interneuron subcategories (all expressing *Reln)*.

We evaluated whether these results were specific to schizophrenia or generally shared across human traits. A heat map of KI Level 1 enrichment *P*-values for GWA results from eight studies of human complex traits are depicted in Figure 2b. Seven of these studies evaluated common variants associations for brain-related diseases or traits with ≥20,000 cases and ≥10 genome-wide significant associations. Human height was included as a non-brain comparator. The results from the earlier PGC GWA study of schizophrenia ^3^ were similar to those from CLOZUK. Although we observed cell types being enriched in other sets none had the specific signal observed in the two schizophrenia sets. For example, major depression disorder (MDD) is another major brain disorder and we found that GABAergic interneurons, embryonic midbrain neurons, and dopaminergic interneurons were the most enriched cell types.

**Figure 2.**
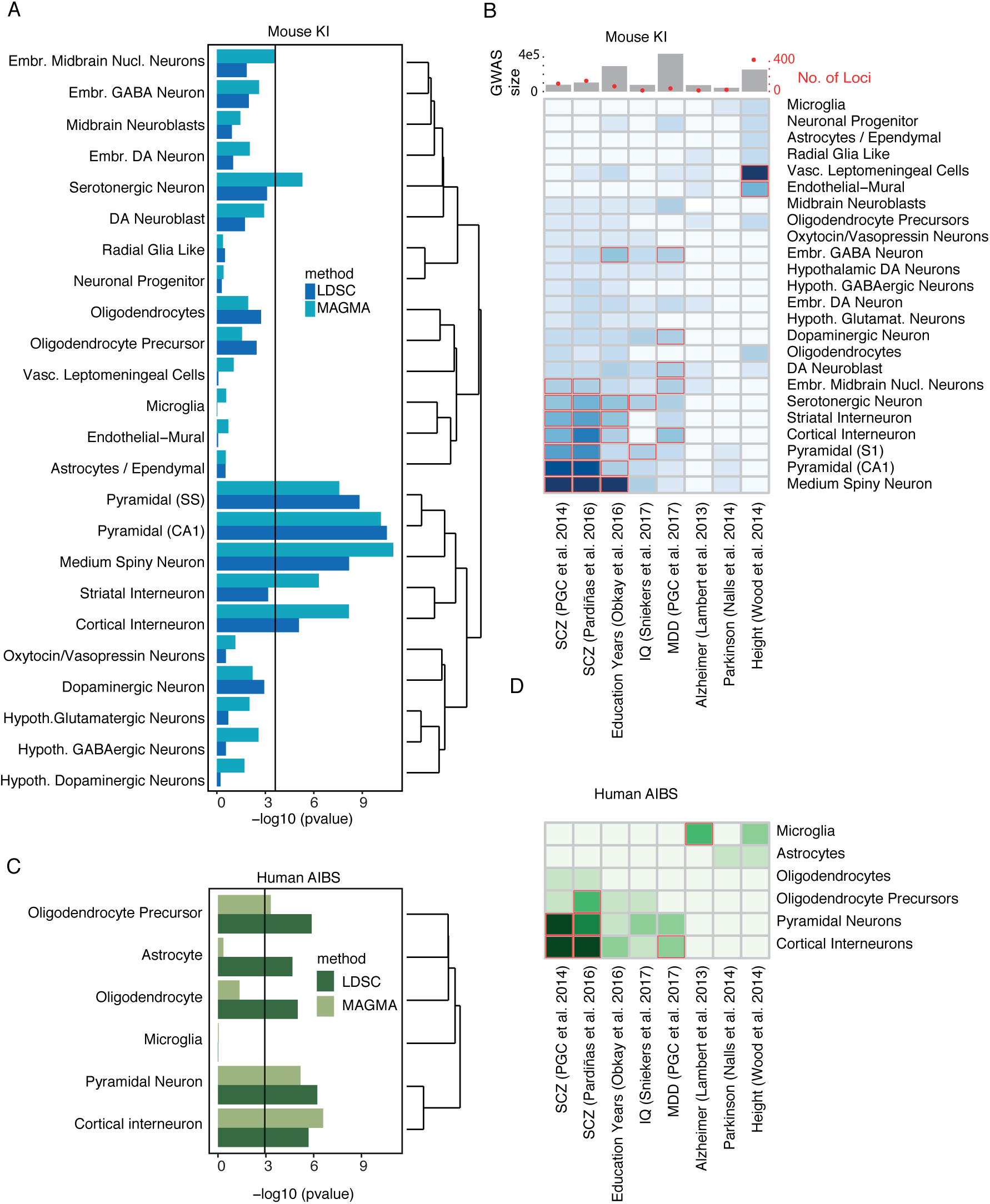
Evaluation of enrichment of common variant CLOZUK schizophrenia GWA results in brain scRNAseq or snRNAseq datasets from adult mouse and human. (A) KI Level 1 brain cell types. Both methods show enrichment for pyramidal neurons (somatosensory cortex and hippocampus CA1), striatal medium spiny neurons, and cortical interneurons. The black line is the Bonferroni significance threshold. (B) Heat map of enrichment probabilities of diverse human GWA with KI Level 1 mouse brain cell types using MAGMA. Bonferroni significant results are marked with red borders. The CLOZUK results do not generalize indiscriminately across human diseases/traits. Unlike schizophrenia, major depressive disorder (MDD) is primarily enriched in cortical interneurons and embryonic midbrain neurons. Total number of cases and controls used in the GWAS are shown in the top bar plot where red dots mark the number of genome-wide significant loci identified. (C) Human mid-temporal cortex Level 1 brain cell types from snRNAseq. Cortical pyramidal neurons and cortical interneurons show significant enrichment. Oligodendrocyte precursors also show enrichment that was not observed in the KI Level 1 data. (D) Heat map of enrichment of diverse human GWA with human mid-temporal cortex Level 1 brain cell types using MAGMA. The CLOZUK results do not generalize across human diseases. MDD again shows significant enrichments in cortical interneurons. Common variant genetic associations for Alzheimer’s disease were enriched in microglia.

Using independent scRNA/snRNAseq mouse brain studies, we replicated our results and found significant enrichment for schizophrenia in hippocampal CA1 pyramidal cells, neocortical pyramidal cells, cortical interneurons, and medium spiny neurons ^19−21^ (Figs. S7a-c). Turning to human data, we evaluated a snRNAseq dataset from mid-temporal cortex (Allen Institute for Brain Science, unpublished), and we confirmed enrichment in cortical pyramidal neurons and cortical interneurons (glutamatergic and GABAergic cells, Figure 2c). The specificity of the temporal cortex signal was confirmed in relation to the same set of brain-specific GWA studies (Figure 2d). Although oligodendrocyte precursors showed up as significant in the human data, it is hard to judge if this is related to a loss of neuronal-specific signal from neurons due to differences in nuclei versus cell sampling (Figure S2). In a small scRNAseq study of human adult and fetal cortex ^22^, adult and fetal cortical neurons were significantly enriched. These are likely pyramidal cells but the small size of this study precluded further exploration (data not shown). No significant enrichments were found in another snRNAseq study of one human ^23^, perhaps due to a lack of cellular diversity (data not shown). We are unaware of scRNAseq data from human hippocampus or striatum. In summary, all major findings from the KI dataset were replicated in independent mouse studies, and the cortical pyramidal cell and cortical interneuron findings were also replicated in independent human studies.

We then evaluated whether gene sets previously implicated in schizophrenia (Table S1) were specifically expressed in the KI level 1 brain cell types using EWCE *(25)*. First, we evaluated pharmacologically-defined molecular targets of antipsychotics (the mainstay of treatment for schizophrenia), which were also previously associated with schizophrenia ^24^. As shown in Figure 3a, antipsychotic medication targets were associated with the same cell types that we found using the CLOZUK GWA schizophrenia results: neocortical S1 pyramidal cells, medium spiny neurons from the dorsal striatum, and hippocampal CA1 pyramidal cells, while cortical interneurons were just above the significance threshold after multiple testing comparison.

**Figure 3.**
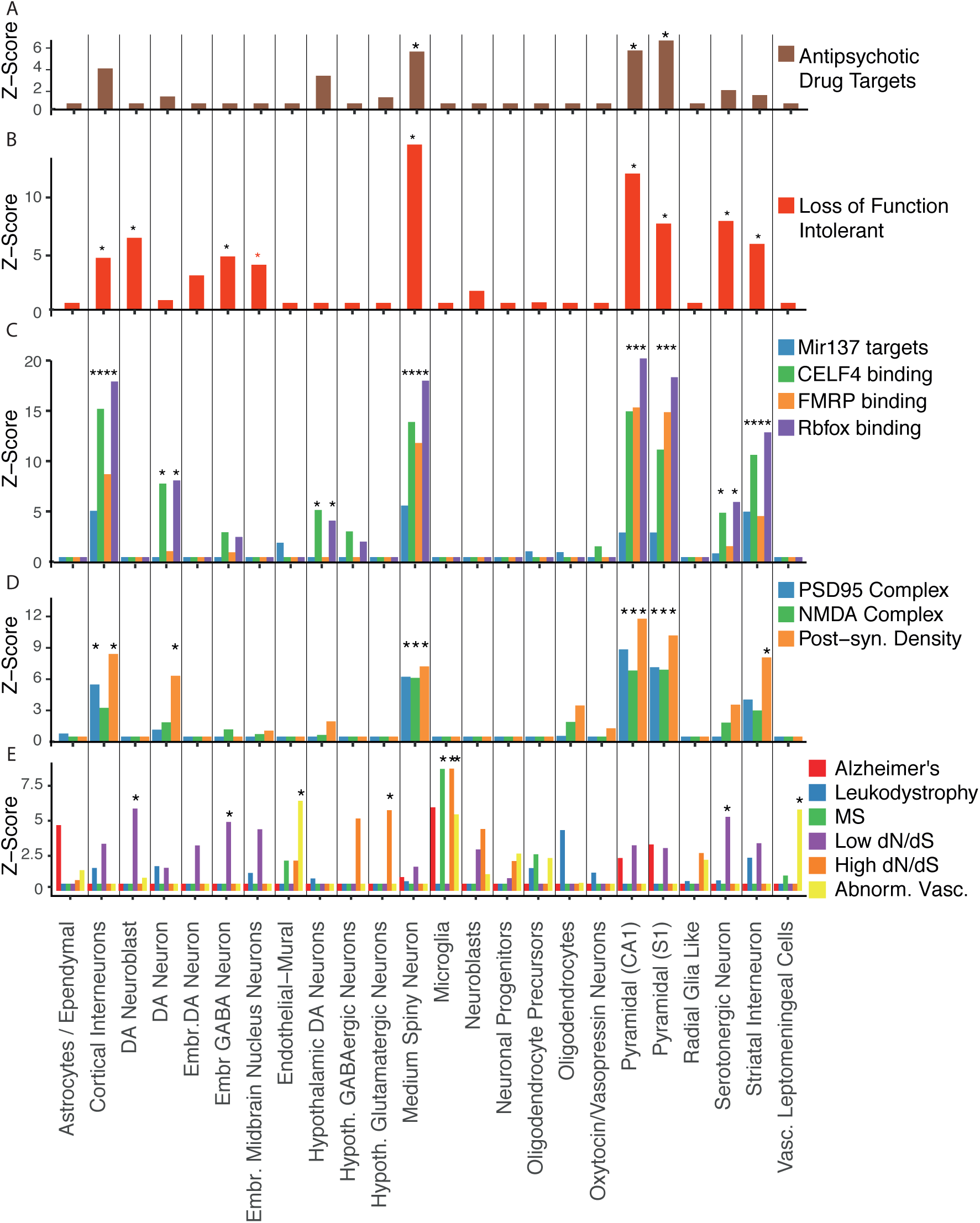
Cell class enrichment of schizophrenia-associated gene sets (Table S1) using EWCE. (A) Antipsychotic medication targets. (B-D) Gene sets previously shown to be enriched for schizophrenia risk genes (B) Genes intolerant to loss-of-function variation. (C) Gene sets defined by known DNA or RNA interactions. (D) Synaptic gene sets. (E) Gene sets unrelated to schizophrenia: Alzheimer’s disease susceptibility genes; genes associated with Human Phenotype Ontology terms for leukodystrophy; multiple sclerosis susceptibility genes and abnormal vasculature; and the top 500 genes with lowest or highest dN/dS ratios between human and mice (i.e., non-synonymous to synonymous exon changes, a measure of conservation). Nearly all KI Level 1 cell types associated with schizophrenia (medium spiny neurons, pyramidal CA1, pyramidal SS, and cortical interneurons) show enrichment for gene sets in A-D. The gene sets shown in E show distinct cell type enrichments which largely correspond to prior expectations. Asterisks denote Bonferroni corrected p-value <0.05.

We then investigated whether other gene sets previously associated with schizophrenia were specifically expressed in schizophrenia relevant cell types (Figure 3b–d). The gene sets consistently associated with schizophrenia – intolerant to loss-of-function variation, NMDA receptor complex, post-synaptic density, RBFOX binding, CELF4 binding, and FMRP associated genes – all had more specific expression in neocortical S1 and hippocampal CA1 pyramidal cells, medium spiny neurons from the dorsal striatum, and cortical interneurons (6 out of 7 but not NMDA receptor complex). These gene sets are involved in diverse cellular functions and, as expected, some gene sets were associated with different KI Level 1 cell types. For example, genes intolerant to loss-of-function variation had greater expression in progenitor cells (dopaminergic neuroblasts, neuroblasts, and embryonic GABAergic neurons).

To evaluate whether these enrichments could be due to some inherent structure of the data, we investigated enrichment of gene sets previously associated with glial cells (Alzheimer’s disease and multiple sclerosis susceptibility genes), associated with Mendelian disorders with clear cellular origins (leukodystrophy and abnormal vasculature), or had either weak or strong conservation (low or high dN/dS scores between humans and mice). None of the enrichment profiles looked similar to that of the schizophrenia-associated lists.

We next assessed how much of the cell type-specific schizophrenia heritability was from overlapping gene expression between cell types. For instance, the association of cortical interneurons is weaker than that of MSNs, but both are GABAergic neurons, and conceivably enrichment in cortical interneurons could be detected due to shared expression of disease genes. This hypothesis can be tested using a resampling without replacement: if the interneuron enrichment is driven solely by overlapping genes with MSNs, then an equivalent level of interneuron association should be obtainable if the scores of genes within each MSN specificity decile are randomized (Figure S8). We performed this randomization 10,000 times for each of the KI level 1 cell types while controlling for all four of the significantly associated cell types (Figure 4a). This analysis revealed that the enrichments of MSNs, cortical interneurons, and hippocampal CA1 pyramidal neurons are independently associated with schizophrenia (due to non-overlapping gene expression). The association with somatosensory pyramidal neurons was found to be largely from genes also expressed by hippocampal CA1 pyramidal neurons.

**Figure 4.**
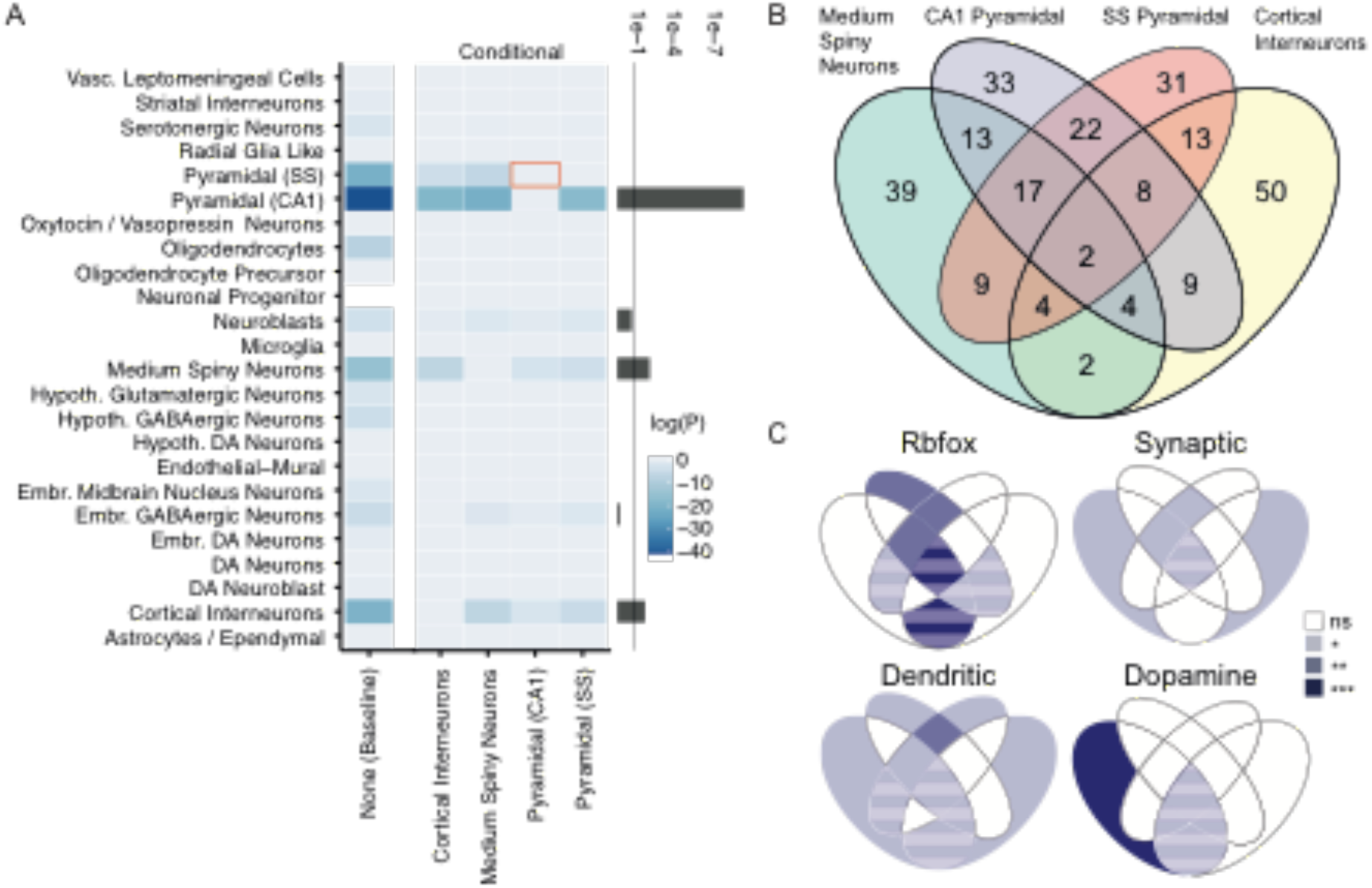
CA1 pyramidal neurons, medium spiny neurons, and cortical Interneurons are independently associated with schizophrenia and distinct molecular pathways contribute to each cell type. (A) Conditional enrichment analysis accounting for correlated gene expression between cell types. The leftmost column show baseline cell type enrichment probabilities values P_celltypeY,baseline_ for schizophrenia calculated by directly fitting a linear model to specificity deciles against MAGMA gene enrichment z-scores (where cell type Y denotes the cell type labelled on the y-axis). The central four columns show P_celltypeY,celltypeX_, the enrichment probability of cell type Y after accounting for correlated expression in cell type (calculated using a resampling method described in the methods). Values of log(P_celltypeY,celltypeX_) approaching zero indicate that after accounting for expression of cell type X, there is no enrichment in cell type Y. The red box highlights that after accounting for expression in CA1 pyramidal neurons there is no longer any enrichment in somatosensory pyramidal neurons; note that the converse is not true. The bar plot on the right shows the minimum value of P_celltypeY,celltypeX_ excluding self-self-comparisons; the vertical line marks p=0.05. (B) Overlap of genes in the schizophrenia-associated cell types. Venn-diagram of the top 1,000 schizophrenia-associated genes from the highest enrichment-deciles in the four Level 1 cell types. (C) P-values for enrichment of genes in the Figure 4b Venn diagram regions. There was enrichment for Rbfox in CA1 pyramidal cells and dopamine signaling in medium spiny neurons along with shared synaptic genes between pyramidal cells but separate for GABAergic cells. Areas with striped shading indicates region with gene number <10.

To use a more qualitative measure of whether heritability associated with enriched cell types was due to shared or distinct sets of genes we plotted the overlap of the top 1,000 genes associated with schizophrenia (via MAGMA) in the 10% most specific genes of each enriched class (Figure 4b). Approximately half of the schizophrenia-associated genes enriched in pyramidal cells and medium spiny neurons were shared but those conferring risk-enrichment in interneurons were to a larger extent exclusive. We then evaluated enrichment of selected gene sets (Rbfox binding genes, genes involved in synapse function, dendritically transported genes, and genes involved in dopaminergic signaling; ***Supplementary Methods***) in the different areas of Figure 4b using a hypergeometric test. The most associated Rbfox genes were enriched in CA1 pyramidal cells, genes related to dopamine signaling were enriched in medium spiny neurons, and synaptic genes associated with schizophrenia were shared between CA1 and S1 pyramidal cells but largely separate in cortical interneurons and medium spiny neurons (Figure 4c). These findings show that each larger neuronal class express a non-overlapping set of risk genes even within the same functional set (e.g. synapse). Our results provide a parsimonious explanation: the primary genetic results pointed at a limited set of brain cells, and the gene sets associated with schizophrenia (including antipsychotic medication targets) pointed at the same cells. The strong enrichment found in the mouse data was at least partly confirmed in the limited human data sets available. Thus, our results suggest that these discrete cell types are central to the etiology of schizophrenia, and provide an empirical rationale for deeper investigation of these cell types in regard to the basis of schizophrenia. These results can be used to guide *in vivo* studies and *in vitro* modeling (e.g., patient-derived iPSCs) and provide a basis for analyzing how different risk genes interact to produce the symptoms of schizophrenia. This approach can be generalized to understand other genetically complex brain disorders.

## Acknowledgements

JHL was funded by the Swedish Research Council (Vetenskapsrådet, award 2014–3863), StratNeuro, the Wellcome Trust (108726/Z/15/Z), and the Swedish Brain Foundation (Hjärnfonden). PFS gratefully acknowledges support from the Swedish Research Council (Vetenskapsrådet, award D0886501). NS was supported by the Wellcome Trust (108726/Z/15/Z). JB was supported by the Swiss National Science Foundation. The PGC has received major funding from the US National Institute of Mental Health (U01 MH109528 and U01 MH109532).

## Materials and Methods

## Online Methods

### Rationale

The overall goal of this analysis was to attempt to connect human genomic findings to specific brain cell types defined by their gene expression profiles: to what specific brain cell types do the common variant genetic findings for schizophrenia best “fit”? Multiple studies have approached this issue ^3,9,13^, but using gene expression based on aggregates of millions of cells. We also evaluated whether gene sets previously implicated in schizophrenia mapped to similar or different brain cell types. We focused on the KI scRNAseq data from mouse (Figure 1 and Table S2). We did this because:

a. These comprise the largest dataset currently available generated using identical procedures. As shown in Figure 1, the total cells with scRNAseq (9,970) and Level 2 cell types (149) exceed all other studies.
b. The mouse data include more brain regions than in human. These regions include a better sampling of those believed to be important in schizophrenia (e.g., currently no data from human striatum or adult dopaminergic neurons).
c. Due to the use of unique molecular identifiers in the KI data, the scRNAseq data reflect absolute counts, and are directly comparable across experiments (particularly for our goal of evaluating enrichment).
d. The mouse data appear to have better signal quality. This could be due to better experimental control or the ability to isolate whole cells (excluding distal neurites) of good quality from mouse but only nuclei or lower quality cells from adult humans. For example, sampling 1,500–3,000 cells in cortical mouse data sets (KI and Tasic et al. ^21^) allowed identification of 24 and 42 cortical neuronal subtypes. In contrast, sequencing over 3,000 human neuronal nuclei^23^ or 466 whole neurons ^22^ allowed for the identification of only 16 and 7 subtypes. More types of inhibitory interneurons (16–23) have been identified in mouse but only 8 in human despite equal or greater sequencing depth but future work may improve the ability to discriminate cell types using single nuclei RNA-Seq data.
e. Use of laboratory mice allow far greater experimental control of impactful perimortem and postmortem events. All mice are healthy without systemic illnesses and medication-free. All mice can be euthanized in the same way, and time from death to tissue processing is standardized and measured in minutes rather than hours. Causes of death in human are highly variable, and perimortem events can alter brain gene expression (e.g., systemic disease or prolonged hypoxia). Although human brain tissue can be obtained during certain neurosurgical procedures (e.g., resection of a seizure focus in refractory epilepsy), the individuals undergoing these procedures are atypical and subject to the effects of chronic brain disease and medication.

Thus, from a practical perspective, more cell types have been identified in mouse, and the KI data comprise over half of currently available brain scRNAseq data. Key findings in the KI dataset were verified in other human and mouse datasets. We also applied independent statistical methods predicated on different assumptions and algorithms to evaluate the relation of brain cell types to GWA results for schizophrenia.

### Limitations

Nonetheless, despite our use of multiple statistical methods and efforts to identify and resolve any spurious explanations for our findings, our work has to be considered in light of inevitable limitations. <u>First</u>, although we used the largest available schizophrenia GWA dataset, we still have an incomplete portrait of the genetic architecture of schizophrenia. This is an active area, and more informative results are sure to emerge in the next few years. <u>Second</u>, although the KI scRNAseq data cover a broad range of brain regions thought to be relevant to the neurobiology of schizophrenia, extensive coverage of cortical and striatal development is lacking at present (gestation, early postnatal, or adolescence).

<u>Third</u>, we focus principally on mouse scRNAseq data. Our reasons for doing so are explained above. A key part of our approach is replication of the main findings in human datasets. However, we would be remiss not to consider the comparability of mouse and human. Mice are widely used for modeling brain diseases. There are relatively high degrees of mouse-human conservation in genes expressed in brain. The most extensive study compared RNA-seq data from six organs (cortex, cerebellum, heart, kidney, liver, and testis) across ten species (human, chimpanzee, bonobo, gorilla, orangutan, macaque, mouse, opossum, platypus and chicken) ^25^. Using principal components analysis, the largest amount of variation (PCs 1 and 2) explained differences between organs rather than between species. Gene expression in brain (including several key gene expression modules) was more conserved between species than any of the other tissues. These observations were broadly replicated using scRNAseq in ventral midbrain ^26^. Furthermore, 75% of genes show similar laminar patterning in mouse and human cortex^27^.

<u>Fourth</u>, whatever the general similarities, there are certainly differences between mouse and human brain ^26^, and there are even cortical cells present in human but not mouse (e.g., spindle or von Economo neurons). ***We therefore evaluated mouse-human gene conservation*.** Using empirical measures of gene conservation (Ensembl, URLs), we determined that the mouse genes in the KI Level 1 and Level 2 gene expression dataset that we analyzed were 89% identical (median, interquartile range 80–95%) to human 1:1 homologues. For these genes, the ratio of non-synonymous to synonymous amino acid changes (dN/dS) was 0.094 (median, interquartile range 0.045–0.173): mutations in these genes are thus subject to strong negative selection (dN/dS = 1 is consistent with neutrality). Pathway analysis of the 400 genes with the largest dN/dS values revealed enrichments in genes involved in defense responses, inflammation, cytokines, and immunoglobulin production. The 400 genes with extremely low dN/dS ratios were involved in neuron differentiation, RNA splicing, and mRNA processing.

In conclusion, for most brain cell types, use of KI mouse scRNAseq data was defensible and reasonable (particularly given verification in human transcriptomic data). The major caution is with respect to cells with prominent immune function (e.g., microglia). (See also the section on mouse-human gene mapping below.)

*We can implicate a particular cell type (i.e., present consistent positive evidence) but it is premature to exclude cell types for which we do not have data, or those with dissimilar function or under selection pressure between mouse and human*.

### Single-cell transcriptome data

Table S2 shows the single nuclei or scRNAseq data from adult mouse or human brain. These include published and unpublished data (using the same protocols as in peer-reviewed papers). To the best of our knowledge, these comprise all or nearly all of the available adult brain single nuclei or scRNAseq data. Most of the available data are from mouse, and a large fraction of the human data are from one person ^23^.

We focused on a superset of brain scRNAseq data from KI generated using identical methods from the same labs with the use of unique molecular identifiers that allow for direct comparison of transcription data across regions (see above for full rationale). The KI mouse superset of 9,970 cells and 149 Level 2 cell types is more extensive than any other single nuclei or scRNAseq dataset now available, and includes most brain regions thought to be salient to schizophrenia. The papers contain full method details. Briefly, the KI scRNAseq data were generated using the same methods (Fluidigm C1 with Illumina 50 bp single end sequencing) with the use of unique molecular identifiers to enable absolute molecular counts. In the first paper describing the method it was estimated that an average of 1.2 million mapped reads per cells was sequenced *(28)*. Level 1 and 2 clustering was done using the BACKSPIN algorithm ^28^. All cells lacking annotations were excluded. For non-neuronal populations, except cells from oligodendrocyte lineage and VLMCs, we only included cells from Zeisel et al 2015 in the KI data set. The level 2 CA1 pyramidal cell contain a small number of cells from CA2 and Subiculum resulting from dissection inaccuracies, these are represented as separate level 2 classes. The resulting data have been shown to be insensitive to linear variation in total reads per cell. If a gene was detected in one dataset and not in another, it was considered to have zero reads in all cells where it had not been detected.

To confirm that no batch effects exist across KI regional subdatasets that may influence the merged results, we plotted three cell types using tSNE which were expected to show little real regional variation: endothelial cells, vascular smooth muscle cells and microglia (Figure S9). The tSNE plots were generated in R using the Rtsne and Scater packages using 500 of the most variable features. Only the embryonic midbrain cells clustered separately, as was expected due to the difference between the embryonic and adult brain.

We include unpublished data generated by the Hjerling-Leffler and Linnarsson labs at KI using the same methods as in Zeisel et al. ^28^. Cells were isolated from dorsolateral striatum from p21-p30 transgenic mice, the same age span as in Zeisel et al. ^28^. Coverage of rare interneuron populations was enhanced by FACS sorting cells from either 5HT3a-EGFP or a Lhx6cre::TdTomato line. The cortical parvalbuminergic cells and striatal neurons were captured and prepared for sequencing as described in Zeisel et al.^28^.

The largest human dataset is an unpublished data set from the Allen Institute for Brain Science which consisted of 4401 cells from middle temporal gyrus of 3 post-mortem brains from healthy, adult subjects. Nuclei were dissociated from cortical tissue and FACS isolated based on NeuN staining, resulting in approximately 90% NeuN+ and 10% NeuN- nuclei. Single nucleus cDNA libraries were generated using SMARTerV4 and Nextera XT and sequenced to a depth of approximately 2 million reads per sample. Reads were aligned with Bowtie and gene expression quantified with RSEM plus intronic reads and normalized to counts per million. Clustering was performed with iterative PCA and tSNE with cluster robustness assessed with 100 bootstrap replicates. Level 1 clusters were characterized based on expression of known marker genes and included two broad classes of neurons –GABAergic interneurons and glutamatergic projection neurons – and 4 non-neuronal types: astrocytes, oligodendrocyte precursors, mature oligodendrocytes, and microglia.

### Mouse-to-human gene mapping

Because most of the scRNAseq data were from mouse brain and the schizophrenia genomic results are from human, it was necessary to map 1:1 homologs between *M. musculus* and *H. sapiens*. To accomplish this, used a best-practice approach in consultation with a senior mouse geneticist (UNC Prof Fernando Pardo-Manuel de Villena de L’Epine, personal communication). We used the expert curated human-mouse homolog list (Mouse Genome Informatics, Jackson Laboratory, URLs, version of 11/22/2016). Only genes with a high-confidence, 1:1 mapping were retained. A large fraction of non-matches are reasonable given evolutionary differences between human and mouse (e.g., the distinctiveness of olfactory or volmeronasal receptor genes given the greater importance of smell in mouse). Nonetheless, we evaluated the quality and coherence of the mapping.

- The mouse brain cell expression levels for the KI Level 1 cell types were similar for mouse genes with and without a 1:1 human homologue. This is inconsistent with a strong bias due to the success/failure of identifying a human homologue.
- A high fraction (93%) of the KI genes detected in mouse brain samples that mapped to a human gene were expressed in human brain (CommonMind DLPFC RNA-seq) or 27 samples with RNA-seq from the Sullivan lab (unpublished, DLPFC from 9 schizophrenia cases and 9 controls plus 9 fetal frontal cortex samples). The ones that did not (7%) were expressed at considerably lower levels in mouse brain or in cell types not prevalent in cortex.
- Of genes with evidence of expression in human brain (via frontal cortex RNA-seq as noted above), human homologues of KI mouse genes accounted for 93.2% of intellectual disability genes, 93.7% of developmental delay genes, 93.8% of genes with a CHD8 binding site, 94.4% of post-synaptic genes, 95.0% of proteins involved in the ciliary proteome, 95.1% of genes intolerant to loss-of-function variation (ExAC pLI > 0.9), 95.6% of pre-synaptic genes, and 96.4% of FMRP interactors.

We evaluated the mapping carefully and the results above suggest the coherence of the mouse-human mapping. All key findings from the KI mouse scRNAseq data were evaluated in other mouse and human brain scRNAseq datasets.

### Calculation of cell type expression proportion

A key metric used for our cell type analyses is the proportion of expression for a given gene. This metric is calculated separately for each single cell dataset (although only one specificity measure is used for the merged KI superset). This is a measure of cell type specificity scaled so that a value of 1 implies that the gene is completely specific to a cell type and a value of 0 implies the gene is not expressed in that cell type. We denote this specificity metric as *s_g,c_* for gene *g* and cell type *c*. Values of *s_g,c_* were calculated for the brain scRNAseq datasets in Table S2.

Each dataset contains scRNAseq results from *w* cells associated with *k* cell types. Each of the *k* cell types is associated with a numerical index from the set {1*,Δ,k}*. The cell type annotations for cell *I* are stored using a numerical index in *L*, such that *l*_1005_=5 indicates that the 1005^th^ cell is of the 5^th^ cell type. We denote *N_c_* as the number of cells from the cell-type indexed by *c*. The expression proportion for gene *g* and cell type *c* (where *r_g,i_* is the expression of gene *g* in cell *i*) is given by:

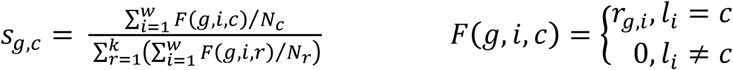

This metric for cell specificity is closely related to other measures ^29^. For instance, the maximum value of *s* per gene yields similar results to *τ* such that *s_max_ >* 0.5 is equivalent to *τ* > 0.94.

### Thresholding of low expressed transcripts

Because *s_g,c_* is independent of the overall expression level of a gene, it is desirable to exclude genes with very low or sporadic gene expression levels, as a small number of reads in one cell can falsely make that gene appear to be a highly specific cell marker. Direct thresholding of low expressed genes is not ideal for performing this as thresholds need to be set individually for each dataset, and some individual cells can show exceptionally and anomalously high expression of the sporadically expressed gene. We reasoned that all the genes we want to include in the study should be differentially expressed in at least one Level 2 cell type included in the study. We thus excluded sporadically expressed genes via ANOVA with the Level 2 cell type annotations as groups, and excluding all genes with *P >* 0.00001. Gene filtering was performed separately for each single cell dataset; importantly though, the KI dataset was filtered as a merged superset. A consequence of this (and of differences in sample preparation and sequencing) is that different genes are used for example in the analysis of the KI superset than were used for the Habib et al (Mouse Hippocampus Div-Seq) dataset. For datasets where level 2 cell type annotations were not available (e.g. the Allan Brain Institute Human Cortex dataset) we used the same approach but with level 1 cell type annotations instead.

### LD Score Regression (LDSC) and partitioning heritability

To partition heritability using LDSC (URLs)^9^, it is necessary to pass LDSC annotation files (one per chromosome) with a row per SNP and a column for each sub-annotation (1=a SNP is part of that sub-annotation). To map SNPs to genes, we used dbSNP annotations (URLs, build 147 and hg19/NCBI Build 37 coordinates). All SNPs not annotated in this file were given a value of 0 in all sub-annotations. Template annotation files obtained from the LDSC Github repository were used as the basis for all cell type and gene set annotations (”cell_type_group.1*”). Only SNPs present in the template files were used. If an annotation had no SNPs, then 50 random SNPs from the same chromosome were selected as part of the annotation (if no SNPs are selected then the software fails to calculate heritability).

Annotation files were created for each cell type for which we applied partitioned LDSC. Twelve sub-annotations were created for each cell type. The first represented all SNPs which map onto named regions which are not MGI annotated genes or which map onto a gene which does not have a 1:1 mouse:human homolog. The second contained all SNPs which map onto genes not expressed in a cell type. The other 10 sub-annotations are associated with genes with increasing levels of expression specificity for that cell type. To assign these, the deciles of *s_g,c_* were calculated over all values of *g (*separately for each value of *c*) to give ten equal length sets of genes. These are then mapped to SNPs as described above. To partition heritability amongst the gene sets (not the cell types), a single set of annotation files was created with each of the gene sets used as a sub-annotation column.

LDSC was then run using associated data files from phase 3 of the 1000 Genomes Project^30^. We computed LD scores for cell type annotations using a 1 cM window (–ld-wind-cm 1). As recommended (LDSC Github Wiki, URLs), we restricted the analysis to using Hapmap3 SNPs, and, as in the original report^9^, these analyses excluded the major histocompatibility region due to its unusual gene density (second highest in the human genome) and exceptionally high LD (highest in the genome). The LDSC “munge_sumstats.py” script was used to prepare the summary statistics files. The heritability is then partitioned to each sub-annotation. We used LD weights calculated for HapMap3 SNPs, excluding the MHC region, for the regression weights available from the Github page (files in the ‘weights_hm3_no_hla’ folder).

For the LD score files used as independent variables in LD Score regression we used the full baseline model ^9^ and the annotations described above. We used the ‘–overlap-annot’ argument and the minor allele frequency files (‘1000G_Phase3_frq’ folder via the ‘–frqfile-chr’ argument, URLs).

Partitioned LDSC computes the proportion of heritability associated with each annotation column while taking into account all other annotations. Based on the proportion of total SNPs in an annotation, LDSC calculates an enrichment score and an associated enrichment *P*-value (one-tailed as we were only interested in annotations showing enrichments of heritability). All figures showing partitioned LDSC results show *P*-values associated with the enrichment of the most specific decile for each cell type.

### Cell type identification using MAGMA

We used MAGMA (v1.04) ^18^, a leading program for gene set analysis ^31^, to evaluate the association of gene-level schizophrenia association statistics with cell-type specific expression under the hypothesis that, in relevant cell types, genes with greater cell type specificity should be more associated with schizophrenia.

Gene level association statistics were obtained using MAGMA (window size 10 kb upstream and 1.5 kb downstream of each gene – see below for discussion window size) using an approach based on Brown’s method ^32^ (model: *snpwise-unweighted)*. This approach allows to combine *P*-values in the specified windows surrounding each gene into a gene-level pvalue while accounting for LD (computed using the European panel of 1000 Genomes Project Phase 3 ^30^).

The tissue specific expression metric for each gene in each cell type was obtained by dividing the gene expression level in a particular cell type by the sum of the expression of the gene in all cell types (see *s_g,c_*, defined above). The distributions of *s_g,c_* were complex (point mass at zero expression, substantial right-skewing).

For each cell type, we transformed *S* into 41 bins (0=not expressed, 1=below 2.5^th^ percentile, 2=2.5-5^th^ percentile, …, 40=above 97.5^th^ percentile), so that each cell type would be comparable.

MAGMA was then used to test for a positive association (one-sided test) between the binned fractions in each cell type and the gene-level associations (option –gene-covar onesided). For a given mouse or human brain cell type, this tested whether increasing tissue specificity of gene expression is associated with increasing common-variant genetic findings for schizophrenia using information from all genes. By default, the linear regression performed by MAGMA is conditioned on the following covariates: gene size, log(gene size), gene density (representing the relative level of LD between SNPs in that gene) and log(gene density). The model also takes into account gene-gene correlations.

In regard to choice of window size/bin boundaries, MAGMA by default combines *P*-values of SNPs located within gene boundaries (±0 kb). We decided to extend the default window size as a large fraction of trait-associated SNPs are located just outside genes in regions likely to regulate gene expression ^3,33^. One of the authors of MAGMA (Christiaan de Leeuw) advised us to expand the window size by a limited amount in order to keep the ability to distinguish the genetic contribution of genes located in close proximity. Therefore, we set expanded gene boundaries to 10 kb upstream-1.5 kb downstream. We evaluated the effect of different choices of bin size including 35 kb upstream-10 kb downstream (as often used by the PGC ^34^), 150 kb upstream-10 kb downstream, and 150 kb upstream-150 kb downstream (GTEx ^17^ Supplementary Figure 9 from). The results were not substantially altered by window size as the ranking of cell types (KI level 1) were very similar for these different window sizes; if anything ours was a slightly conservative choice.

### Enrichment analyses of gene sets and antipsychotic drug targets

Expression Weighted Cell type Enrichment (EWCE, Bioconductor, URLs) ^35^ was used to test for cell types which show enriched expression of genes associated with particular schizophrenia associated gene sets. These analyses used the same specificity *(S)* values for the KI Level 1 data that were used for the MAGMA and LDSC analyses. EWCE was run with 10,000 bootstrap samples. Enrichment *P*-values were corrected for multiple testing using the Bonferroni method calculated over all cell types and gene lists tested. EWCE returns a z-score assessing standard deviations from the mean. Values < 0 (a depletion of expression) were recoded to zero.

### Evaluation of genomic biases

The algorithms used by LDSC and MAGMA both account for the non-independence introduced by linkage disequilibrium (LD), or the tendency for genomic findings to “cluster” due to strong intercorrelations. LD block size (discrete regions of high correlations between nearby genetic markers) average 15–20 kb in samples of European ancestry, but there are nearly 100 genomic regions with high LD extending over 1 mb (the extended MHC region on human chromosome 6 is the largest and has very high LD over 8 mb). Gene size is an additional consideration for MAGMA (accounting for gene size is a component of the algorithm), particularly as brain-expressed genes are considerably larger than genes not expressed in brain (mean of 80.7 kb vs 31.2 kb). The algorithms used by LDSC and MAGMA have been well-tested, and are widely used. However, it is conceivable that certain edge cases could defeat algorithms that work well for the vast majority of scenarios. An example might be if a large fraction of the genes that influence a brain cell type were located in a region of very high LD. First, brain-expressed genes were slightly more likely to be in a large LD block (≥99^th^ percentile in size across the genome), 12.4% vs 10.2%. In discussion with the developers of LDSC and MAGMA, this should not yield an insuperable bias. Second, by counting the numbers of genes and brain-expressed genes per mb, we found that brain-expressed genes in the human genome were reasonably evenly scattered across the genome *(R*^2^ 0.85), and only 10 of 2,534 1-mb intervals were outliers. Most of these were gene clusters with fewer than expected brain-expressed genes (e.g., a late cornified envelope gene cluster on chr1:152–153 mb, an olfactory gene cluster on chr1:248–249 mb, and a keratin gene cluster on chr17:39–40 mb). Third, in a similar manner, we evaluated the locations of the human 1:1 mapped genes influential to the KI Level 1 classifications and found these to be relatively evenly scattered in the genome. Thus, these potential genomic biases did not appear to present difficulties for our key analyses (that used two independent methods in any event).

### Schizophrenia common variant association results

The schizophrenia GWA results were from the CLOZUK and PGC studies ^3,16^. CLOZUK is the largest currently obtainable GWA for schizophrenia (40,675 cases and 64,643 controls), and the authors identified ~150 genome-wide significant loci. It includes the schizophrenia samples from the earlier PGC paper. The CLOZUK manuscript has been submitted, reviewed, revised, and resubmitted, and a preprint is available in biorXiv (DOI 10.1101/068593). For selected analyses, we also included the PGC schizophrenia results from the *Nature* 2014 report, obtained from the PGC download site (URLs). This paper included 36,989 cases and 113,075 controls, and identified 108 loci associated with schizophrenia. Results from the published PGC and CLOZUK studies were qualitatively similar with the CLOZUK data generally showing increased significance owing to its larger sample size.

### Comparison GWA results for other traits

We included comparisons for a selected set of brain related traits as well as height as a negative control. As power to identify cell types is directly proportional to the sample size of the GWA study, we only included traits with at least 20’000 samples that discovered at least 20 genome-wide significant loci. The GWA results were from the following sources: schizophrenia ^3^ from the PGC; Alzheimer’s disease ^36^; educational attainment^37^; IQ^38^; MDD from the PGC (unpublished); Parkinson’s disease^39^ and height^40^.

### Gene sets associated with schizophrenia

The gene set results for schizophrenia are summarized in Table S1, and the gene sets are included in ***Table S4***. For CELF4 binding genes ^41^, we used genes with iCLIP occupancy > 0.2 from Table S4. For FMRP binding genes ^12^, we used genes from Table S2A. Genes intolerant to loss-of-function variation were from the Exome Aggregation Consortium (pLI > 0.9) ^10^. Genes containing predicted miR-137 target sites were from <u>microrna.org</u> (URLs). NMDA receptor complex genes came from Genes-to-Cognition database entry L00000007 ^42^. The human post-synaptic density gene set was from Table S2 ^43^. The PSD95 complex came from Table S1 using all genes marked with a cross in the ‘PSD-95 Core Complex’ column ^44^. For RBFOX binding, we took all genes with RBFOX2 count > 4 or summed RBFOX1 and RBFOX3 > 12 from Table S1 ^45^. For antipsychotic drug targets, we used a gene list provided by Drs Gerome Breen and Héléna Gaspar as reported in the biorXiv preprint (DOI 10.1101/091264).

### Gene sets not associated with schizophrenia

The gene sets are included in ***Table S4***. For Multiple Sclerosis susceptibility genes we used the top results from the MSGene database (http://msgene.org/TopResults.asp). For Alzheimer’s disease we used the top results from the AlzGene database^46^(http://www.alzgene.org/TopResult.asp) as well as genome wide significant genes reported in an additional GWAS study^47^. For genes associated with leukodystrophy (HP:0002415) and abnormal vasculature (HP:0002597) we used the Human Phenotype Ontology^48^(http://compbio.charite.de/hpoweb). To obtain the genes with the top 500 highest/lowest dN/dS between humans and mice we obtained the dN and dS values through BioMart associated with each ensemble gene ID, mapped these onto HGNC gene symbols (averaging where ensemble and HGNC did not have 1:1 matches).

### Depletion of dendritically enriched transcripts in nuclei datasets

The list of dendritically enriched transcripts were obtained from ^15^ supplementary table 10. This list was produced from pyramidal cells from rat hippocampus and human 1:1 homologs were obtained as described above: we refer to this set of genes as L_dendritic_. To enable direct comparisons between datasets, all datasets were reduced to contain a common core of six level 1 celltypes: pyramidal neurons, interneurons, astrocytes, interneurons, microglia and oligodendrocyte precursors. In the case of the KI dataset, we used S1 Pyramidal neurons rather than CA1 Pyramidal. The specificity metric (denoted above as *s_g,c_*) was recalculated for each dataset using this reduced set of celltypes. Comparisons were then made between datasets (denoted in the graph with the format ‘Dataset X vs Dataset Y’). We denote the mean pyramidal neuron specificity scores for dendritically enriched genes in dataset X as 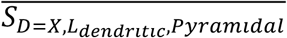. We then get the difference in pyramidal specificity of for list *L* between two datasets as 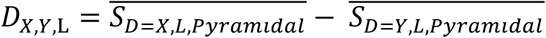. We then calculate values of *D_X,Y,l_* for 10000 random gene lists, having the same length as the dendritically enriched gene list, with the genes randomly selected from the background gene set. We denote the n^th^ random gene list as *R_n_*. The mean and standard deviation of the bootstrapped *D_X,Y,L_* values are denoted *μ_D,Y,R_* and *σ_D,Y,R_* respectively. The depletion z-score is then calculated as: 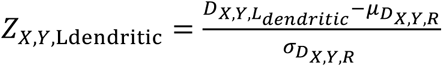. a large positive z-score thus indiciates that dendritically enriched transcripts are specifically depleted from pyramidal neurons from dataset Y relative to dataset X.

### Conditional cell type enrichments

Gene association z-scores for Schizophrenia were calculated in MAGMA as described above. To enable randomisation of the z-scores and recalculation of the associations to be done programmatically, these were then loaded into R and associations with disease were calculated within this environment without external calls to MAGMA. All genes within the extended MHC region (chr6 25-34mb) were dropped from all aspects of this analysis. We controlled for gene size and gene density by regressing out the effect of NSNPS and NDENSITY parameters (and the log of each) on the z-score. To ensure a meaningful number of genes were randomised within each group, associations were calculated over deciles rather than the smaller percentile bins used earlier with MAGMA. Probabilities of association are calculated using the lmFit and ebayes functions from the limma package to enable rapid computation. We denote the set of cells studied as *C* such that *c_i_* represents the i^th^ celltype. The original z-scores are denoted Z such that *z_t_* is the z-score of the i^th^ gene while the randomised z-scores are denoted *R*. The set of genes in the i^th^ specificity decile of the controlled cell type, *c_x_* and the j^th^ specificity decile of target cell type, *c_y_* are denoted 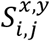 and thus 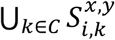 contains all genes in the i^th^ specificity decile of cell cell type, *c_x_*.

The basis of the approach, as depicted in Figure S8 is to randomise the z-scores with respect to the specificity deciles of the target celltype, *c_y_* but not with respect to the specificity deciles of the controlled celltype, *c_x_*. Thus for each of the deciles indexed by i we randomly resampled without replacement the z-scores such that 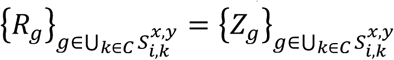 and yet *R_g_ ≠ Z_g_*. In practical terms, this would mean that if we controlled for MSN’s and targeted cortical interneurons, the mean z-score in the 10^th^ MSN decile would remain the same but would be different in cortical interneurons; the question being tested is the degree to which this equates to total randomisation in terms of the schizophrenia association found in cortical interneurons.

The baseline association values shown in Figure 4a leftmost column (described as P_cellty eY,baseline_) were calculated using *Z*. The values of P_celltypeY,celltypeX_ (probability of celltype y being associated with schizophrenia controlling for celltype x) are calculated using intermediate probabilities: 10,000 association p-values are calculated for resampled values of *R*. We selected the 500^th^ lowest of these p-values (equivalent to the value which the baseline association probability would need to exceed to be declared independently associated with a probability of 95%) and denote this 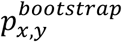. The value of P_celltypeY,celltypeX_ is then calculated as 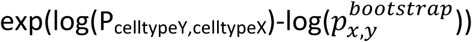. If the value of P_celltypeY,celltypeX_ exceeds 1 (indicating that the randomised samples were actually more significantly associated than was found to be the case) then it is set to 1. We were also able to evaluate whether the probability of schizophrenia association in celltype y is greater than would be expected based solely on the expression in celltype x by asking whether the actual association p-value was lower than 95% of the bootstrapped p-values. As expected, all self-self comparisons were found to be non-significant by this metric (i.e. after accounting for expression in CA1 pyramidal neurons, CA1 pyramidal neurons are no longer significant). In Figure 4a, a red box was placed around the CA1 Pyramidal vs Somatosensory Pyramidal square because this was the only comparison involving the four significantly associated cell types in which controlling for expression of a different cell type abolished the enrichment.

### Venn diagram enrichments

The Venn diagram shown in Figure 4 was generated using by selecting the top 1000 genes most associated with Schizophrenia based on the MAGMA gene specific z-scores. All genes within the extended MHC region (chr6 25-34mb) were dropped from the analysis. We controlled for gene size and gene density by regressing out the effect of NSNPS and NDENSITY parameters (and the log of each) on the z-score. We then took the intersection of the top 1000 genes with the top decile for each of the four significantly associated level 1 cell types and generated the Venn diagram using the R *VennDiagram* package. The dopamine gene set include all genes associated with any of the following GO terms: GO:0090494 (”dopamine uptake”), GO:0090493 (“catecholamine uptake”), GO:0051584 (“regulation of dopamine uptake involved in synaptic transmission”), GO:0032225 (“regulation of synaptic transmission, dopaminergic”), GO:0001963 (“synaptic transmission, dopaminergic”) and GO:0015872 (“dopamine transport”). The synaptic gene list comprised a combination of three published gene lists: the human post-synaptic density (referenced above); presynaptic active vesicle docking sites ^49^ and synaptic vesicle genes ^50^. For the presynaptic gene list, the data came from supplementary table S1, the geneInfo numbers were converted from genInfo accessions to Refseq IDs using Entrez Batch then from Rat RefSeq to HGNC symbols keeping only 1:1 homologs. The synaptic vesicle gene list came from supplementary table S1, and were converted from Rat RefSeq to HGNC symbols using only 1:1 homologs. Enrichment probabilities were calculated using a hypergeometric test against a background set of all MGI genes with 1:1 homologs in human (as described above).

## Supplemental Tables

**Table S1.**
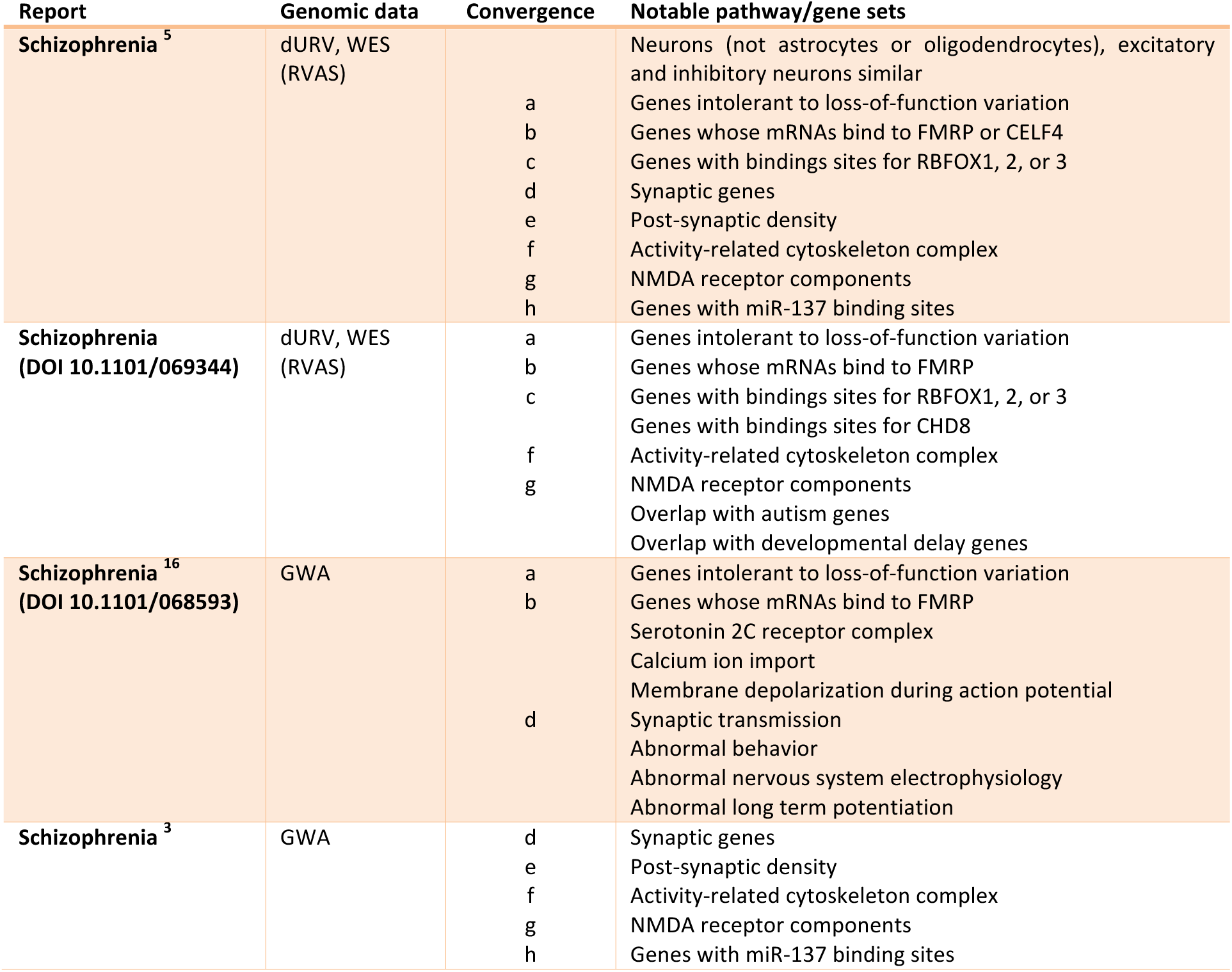
shows the gene sets or biological pathways implicated in schizophrenia. These analyses ask whether schizophrenia case/control genetic association results are “enriched” in the genes comprising a gene set. At least 20 gene sets have been implicated, and many are implicated by different types of genetic studies. This convergence is highly notable. However, these connect genetic risk for schizophrenia to a highly diverse and even puzzling set of genes and biological pathways. dURV=disruptive or damaging ultra-rare variants. WES=whole exome sequencing. GWA=genome-wide, common variant association study. RVAS=rare variant association study.

**Table S2.** Single nuclei or scRNAseq data from mouse or human brain. Source column points to where the data were obtained (URLs). KI=Karolinska Institutet. AIBS=Allen Institute for Brain Science. L1=number of Level 1 cell type categories. L2=number of Level 2 cell types (subdivisions of L1 types). All datasets labelled as KI were merged into a single superset; all other datasets were used separately. These data are depicted in **Figure 1**.

| Species | Label | Citation | Source | RNAseq tech. | Stage | Region | Cells | L1 | L2 |
| --- | --- | --- | --- | --- | --- | --- | --- | --- | --- |
| Mouse | Gokce | <sup>19</sup> | GEO GSE82187 | Single cell | Adult | Striatum | 1208 | 10 | 10 |
|  | Habib | <sup>20</sup> | GEO GSE84371 | Single nuclei | Adult | Hippocampus | 1287 | 11 | 29 |
|  | KI | Unpublished | Pending | Single cell | Adult | Cortex (Pvalb interneurons) | 89 | 1 | 1 |
|  | KI | <sup>28</sup> | Linnarsson lab (URLs) | Single cell | Adult | Cortex + hippocampus | 1996 | 6 | 41 |
|  | KI | <sup>51</sup> | GEO GSE74672 | Single cell | Adult | Hypothalamus | 772 | 4 | 62 |
|  | KI | <sup>26</sup> | GEO GSE76381 | Single cell | Adult | Midbrain | 243 | 1 | 5 |
|  | KI | <sup>26</sup> | GEO GSE76381 | Single cell | Fetal | Midbrain | 1290 | 8 | 22 |
|  | KI | <sup>52</sup> | GEO GSE75330 | Single cell | Adult | Oligodendrocytes | 5051 | 3 | 13 |
|  | KI | Unpublished | Pending | Single cell | Adult | Striatum | 529 | 2 | 6 |
|  | Tasic | <sup>21</sup> | GEO GSE71585 | Single cell | Adult | Cortex | 1679 | 7 | 49 |
| Human | AIBS | Unpublished | Pending | Single nuclei | Adult | Cortex | 4401 | 6 | 6 |
|  | Darmanis | <sup>22</sup> | GEO GSE67835 | Single nuclei | Adult & fetal | Cortex | 420 | 8 | 8 |
|  | Lake | <sup>23</sup> | dbGaP phs000833.v3.p1 | Single nuclei | Adult | Cortex (N=1) | 3042 | 2 | 16 |

**Table S3.**
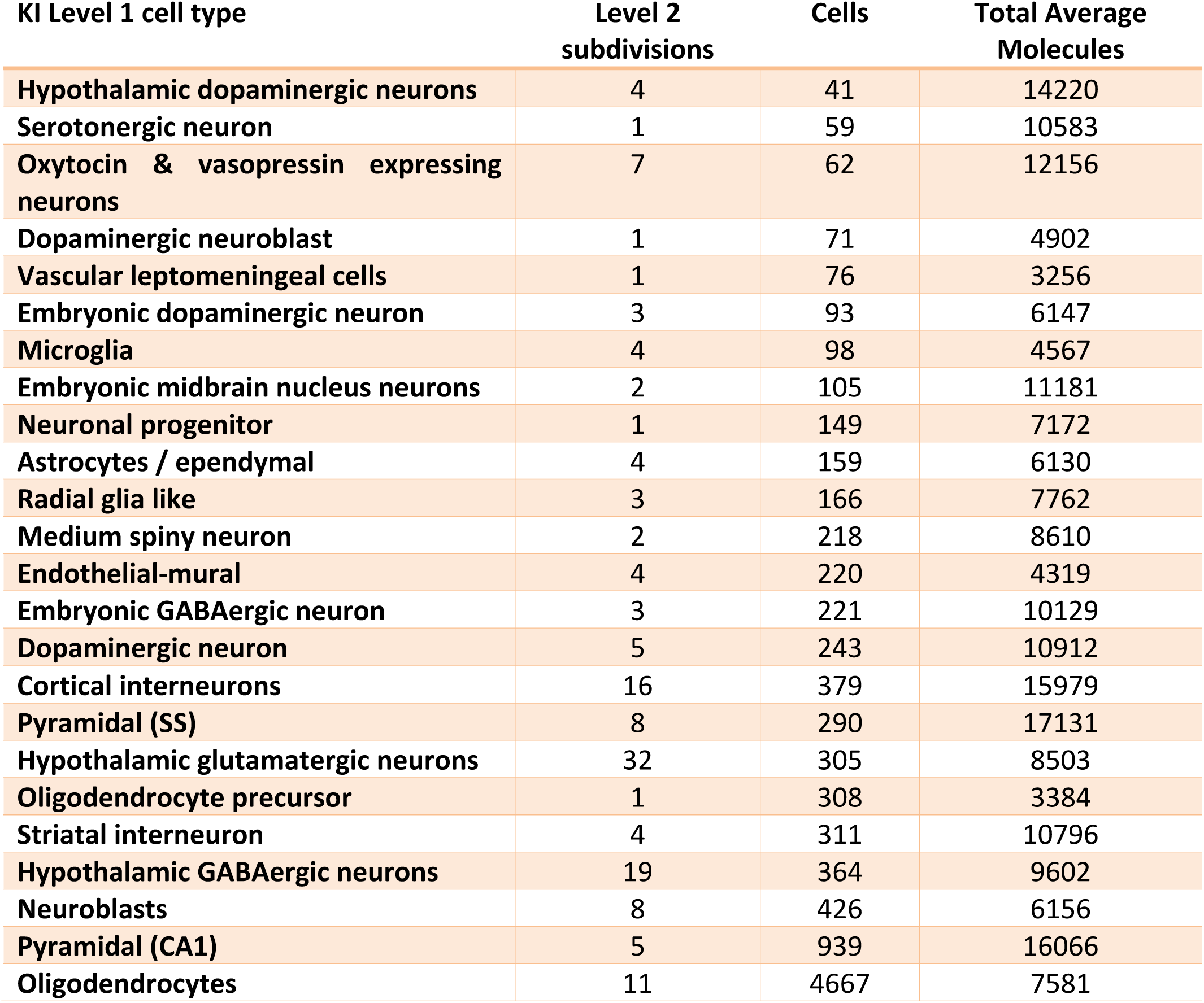
Detail on the KI scRNAseq Level 1 and 2 dataset. There are 24 Level 1 cell type categories and 149 Level 2 subdivisions. The median number of cells in the Level 1 categories was 218 (interquartile range 91–306), and the Level 2 subdivisions range from 1–32. The numbers of single cells contributing to the Level 1 classification are shown. We found no relation between the number of cells and the cell type found to “fit” schizophrenia. For example, there are large numbers of oligodendrocytes and neuroblasts (which were not enriched for schizophrenia genomic findings), and the number of cells for medium spiny neurons (which were associated) were at the median. Likewise, the total average number of molecules detected in each cell type (as determined using Unique Molecular Identifiers) does not explain the enrichments found (note that medium spiny neurons have almost half as many molecules as hypothalamus dopaminergic neurons).

| KI Level 1 cell type | Level 2 subdivisions | Cells | Total Average Molecules |
| --- | --- | --- | --- |
| Hypothalamic dopaminergic neurons | 4 | 41 | 14220 |
| Serotonergic neuron | 1 | 59 | 10583 |
| Oxytocin & vasopressin expressing neurons | 7 | 62 | 12156 |
| Dopaminergic neuroblast | 1 | 71 | 4902 |
| Vascular leptomeningeal cells | 1 | 76 | 3256 |
| Embryonic dopaminergic neuron | 3 | 93 | 6147 |
| Microglia | 4 | 98 | 4567 |
| Embryonic midbrain nucleus neurons | 2 | 105 | 11181 |
| Neuronal progenitor | 1 | 149 | 7172 |
| Astrocytes / ependymal | 4 | 159 | 6130 |
| Radial glia like | 3 | 166 | 7762 |
| Medium spiny neuron | 2 | 218 | 8610 |
| Endothelial-mural | 4 | 220 | 4319 |
| Embryonic GABAergic neuron | 3 | 221 | 10129 |
| Dopaminergic neuron | 5 | 243 | 10912 |
| Cortical interneurons | 16 | 379 | 15979 |
| Pyramidal (SS) | 8 | 290 | 17131 |
| Hypothalamic glutamatergic neurons | 32 | 305 | 8503 |
| Oligodendrocyte precursor | 1 | 308 | 3384 |
| Striatal interneuron | 4 | 311 | 10796 |
| Hypothalamic GABAergic neurons | 19 | 364 | 9602 |
| Neuroblasts | 8 | 426 | 6156 |
| Pyramidal (CA1) | 5 | 939 | 16066 |
| Oligodendrocytes | 11 | 4667 | 7581 |

## Supplemental Figures

**Figure S1.**
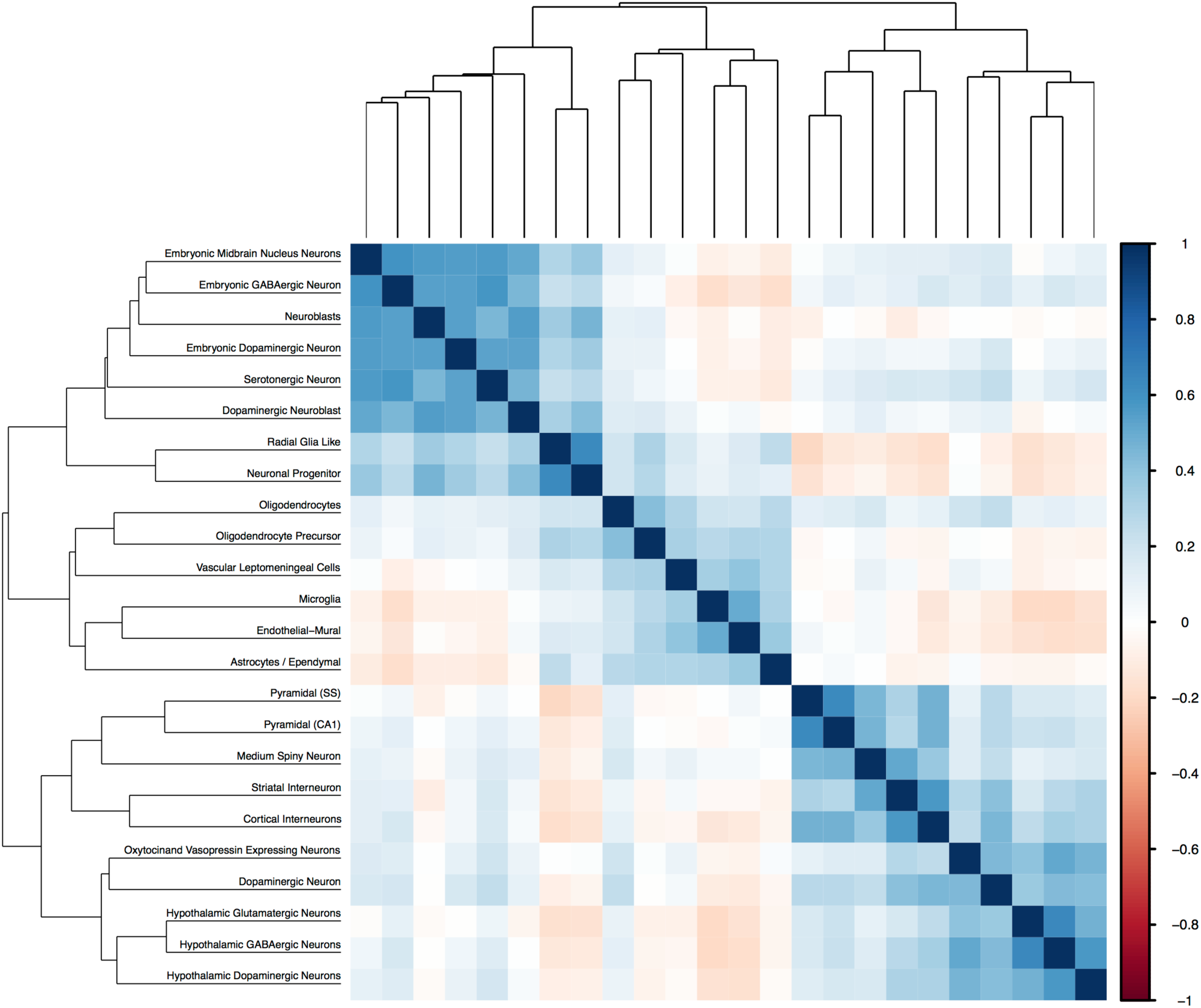
Correlation of the binned measure of cell type-specific expression (S, defined in Online Methods) for mouse brain cell types in the KI level 1 dataset. To illustrate the overall structure of mouse brain scRNAseq dataset, we show a heat map and clustering of the brain cell types identified in the KI Level 1 data. The major divisions are embryonic/progenitor cells (upper left), support cells (middle; e.g., oligodendrocytes and microglia), and mature cells (lower right). The major division of the mature cells include pyramidal cells/medium spiny neurons, interneurons, and “speciality” neurons (i.e., dopaminergic, GABAergic, and glutamatergic).

**Figure S2.**
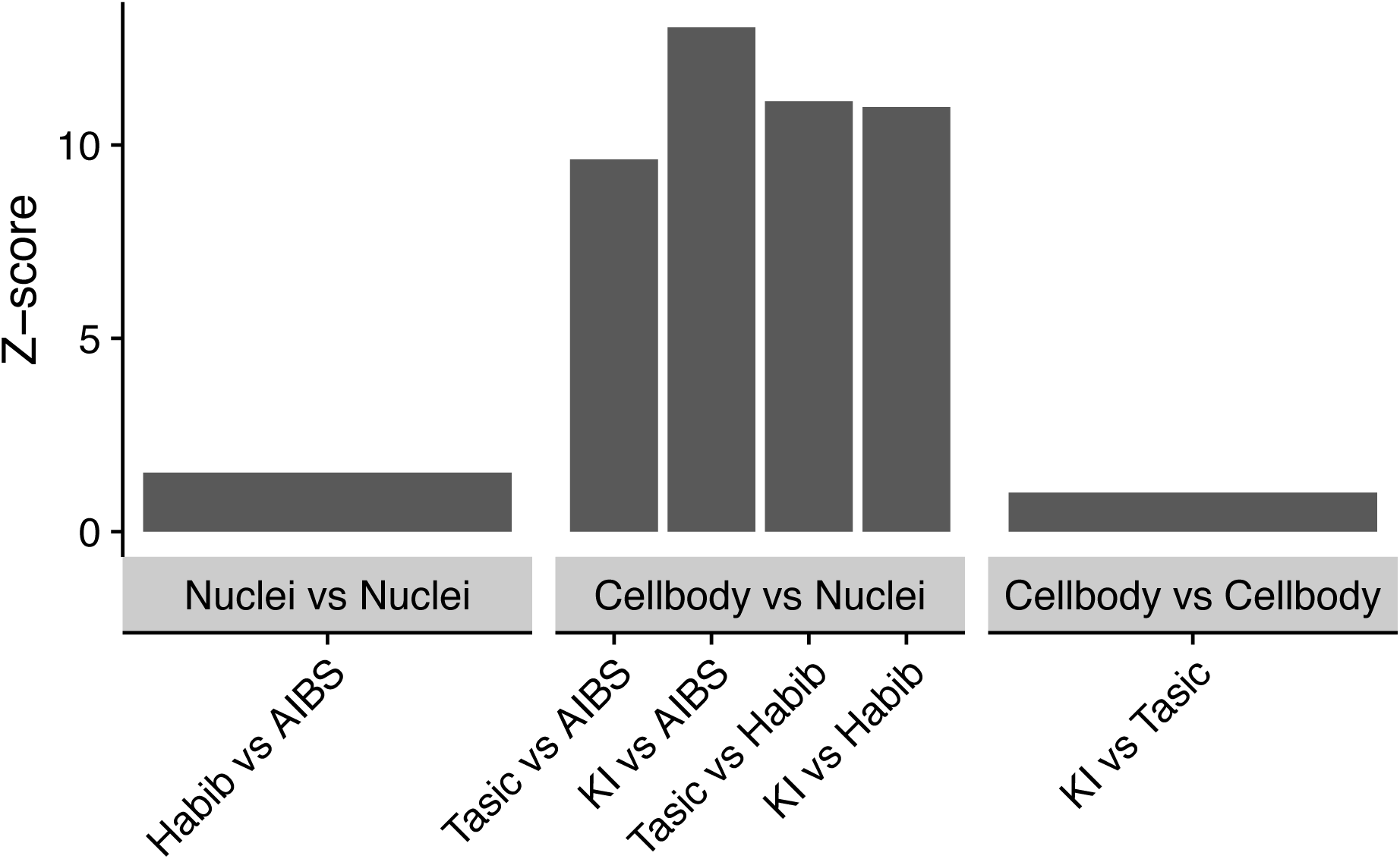
Dendritically enriched transcripts (DETs) are specifically depleted from brain single-nuclei (“nuclei”) RNAseq datasets relative to single-cell (“cell body”) RNAseq data. These 2,252 DETs were identified in a prior study of expression in hippocampal neuropil relative to cell body layer^15^. Each bar represents a comparison between two datasets (dataset X vs dataset Y), with the bootstrapped z-scores representing the extent to which DETs have lower specificity for pyramidal neurons in dataset Y relative to X. Larger z-scores indicate greater depletion of DETs. Table S2 describes the studies. The snRNAseq/nuclei studies were: Habib et al. ^20^ (adult mouse hippocampus) and AIBS (Allen Institute for Brain Science, unpublished, adult human cortex). The scRNAseq/cellbody studies were KI (adult mouse cortex and hippocampus) and Tasic et al. ^21^ (mouse cortex).

**Figure S3.**
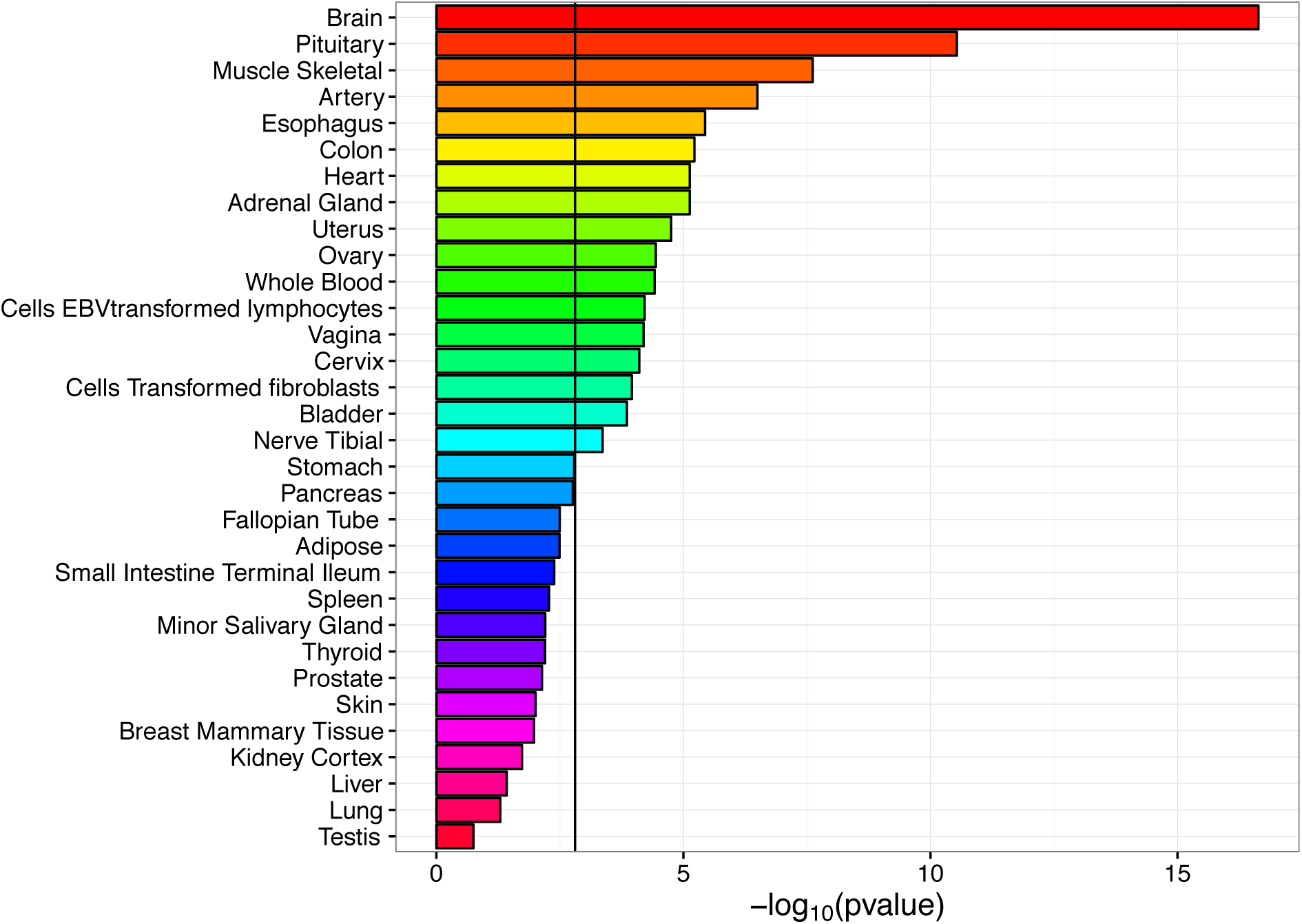
MAGMA enrichment of CLOZUK schizophrenia GWA results in relation to human tissue-specific gene expression (from GTEx). Brain and pituitary are most associated, but there are significant associations with multiple non-brain tissues), and tissues not believed to be etiologically involved in schizophrenia (colon, heart, uterus. Black line shows Bonferroni correction.

**Figure S4.**
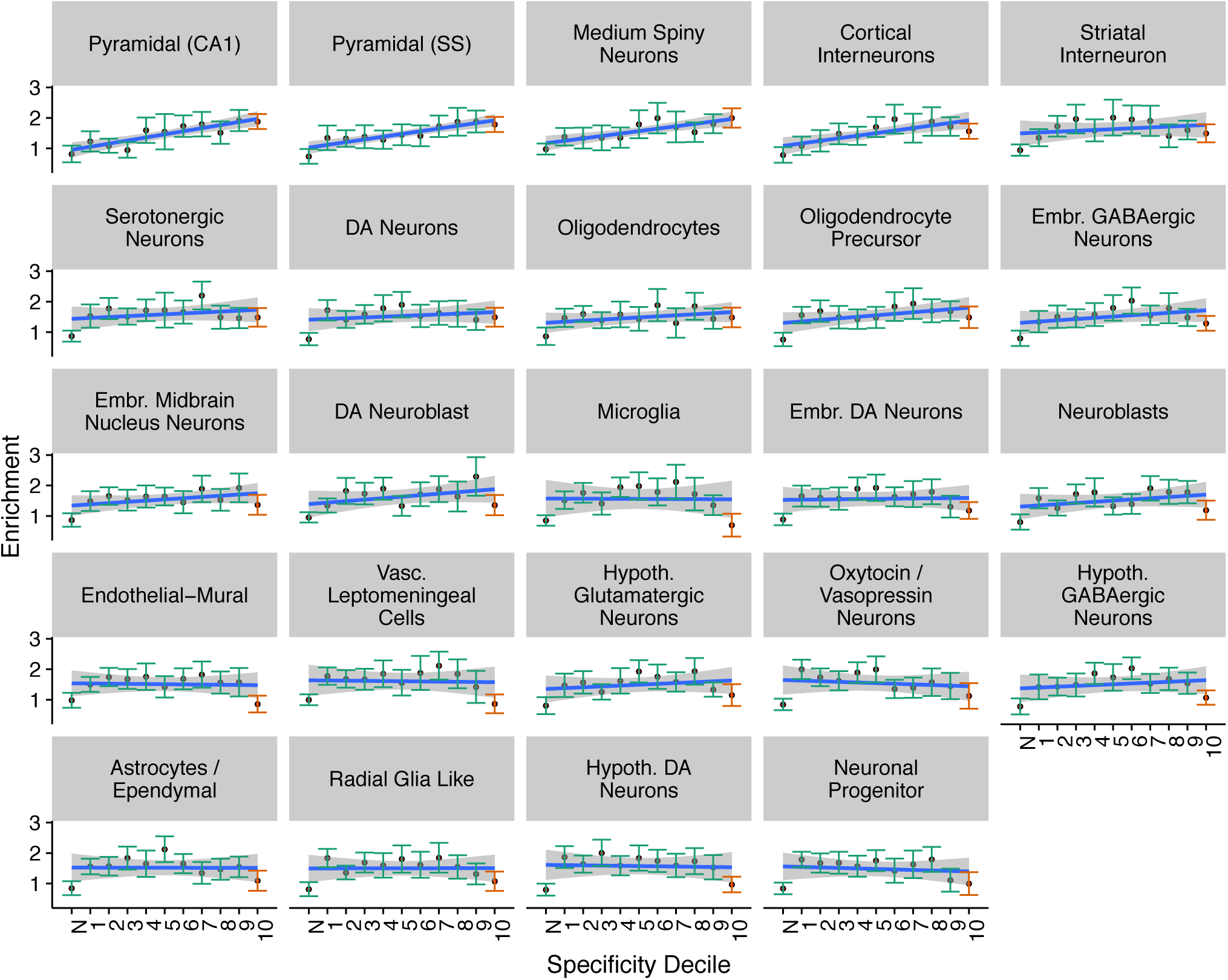
LDSC Schizophrenia (CLOZUK) enrichment values in each specificity decile for each of the KI level 1 cell types. Error bars indicate the 95% confidence intervals. The rightmost point and its confidence intervals are marked in red as this is the decile used for reported LDSC probabilities throughout this paper (rather than the probability of the slope increasing as was used for reporting MAGMA probabilities). The leftmost point (marked ‘N’) represents all SNPs which map onto genes not expressed in MSNs. Blue line slows the linear regression slope fitted to the enrichment values. The grey boxing around the blue regression line depict the confidence intervals of the regression line.

**Figure S5.**
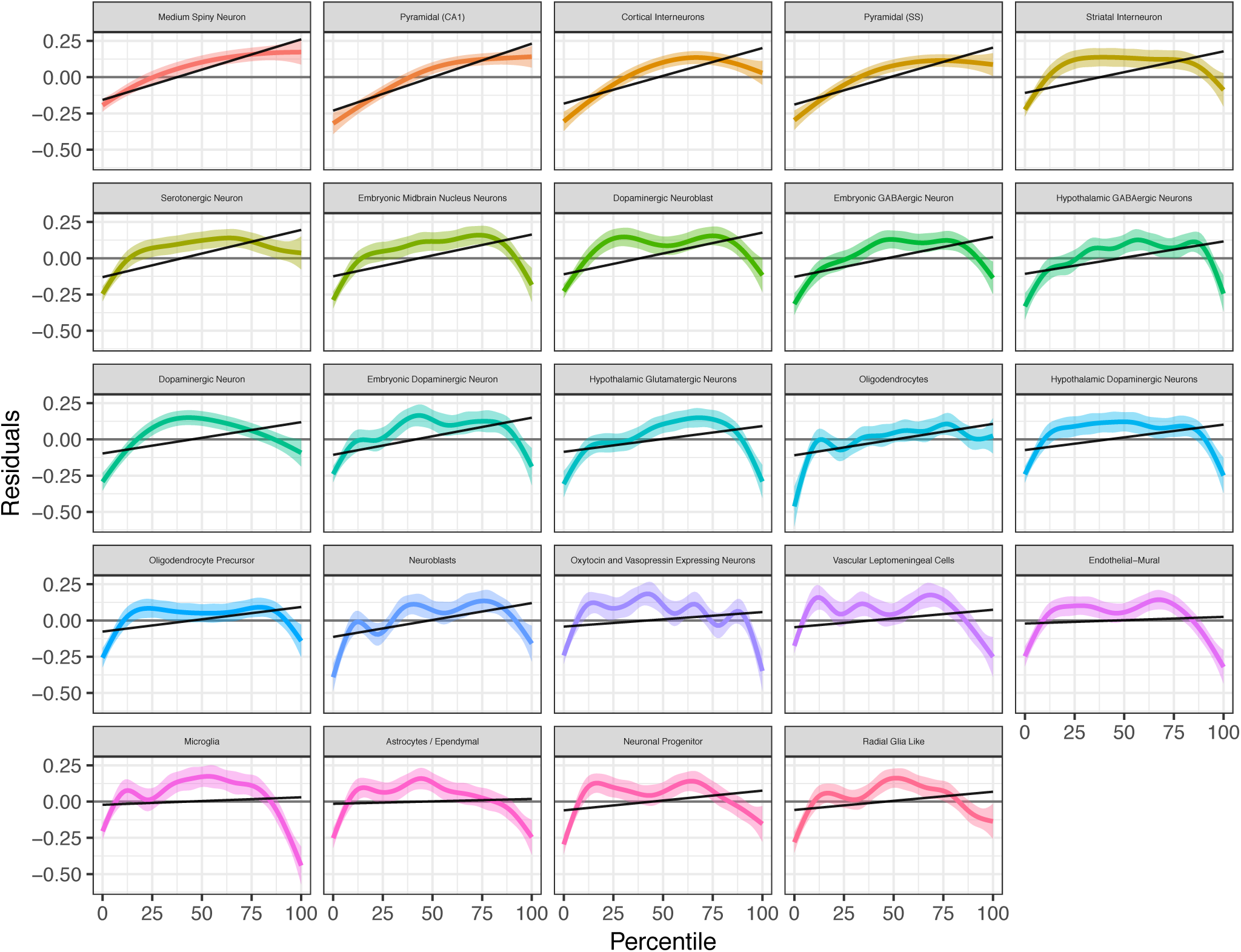
MAGMA Schizophrenia (CLOZUK) gene level model fit for each KI level 1 cell type. The Y-axis (residuals) was obtained by regressing the gene length, gene density and their logs from the gene-level z-score obtained from MAGMA using the CLOZUK schizophrenia GWAS. Negative residuals indicate that genes are less associated with schizophrenia, while positive residuals indicate that genes are more associated with schizophrenia. The x-axis are the 41 binned tissue specificity measures (each bin represent a 2.5% quantile of the distribution of proportions in the cell type) multiplied by 2.5 (the bin 40 which represents genes that are the 2.5% most specifically expressed in the cell type will have a value of 100, etc.). The coloured line shows the best non-linear fit to the data using a generalised additive model (GAM) with its 95% confidence interval. The black line represents the linear regression of the residuals by the binned proportions

**Figure S6.**
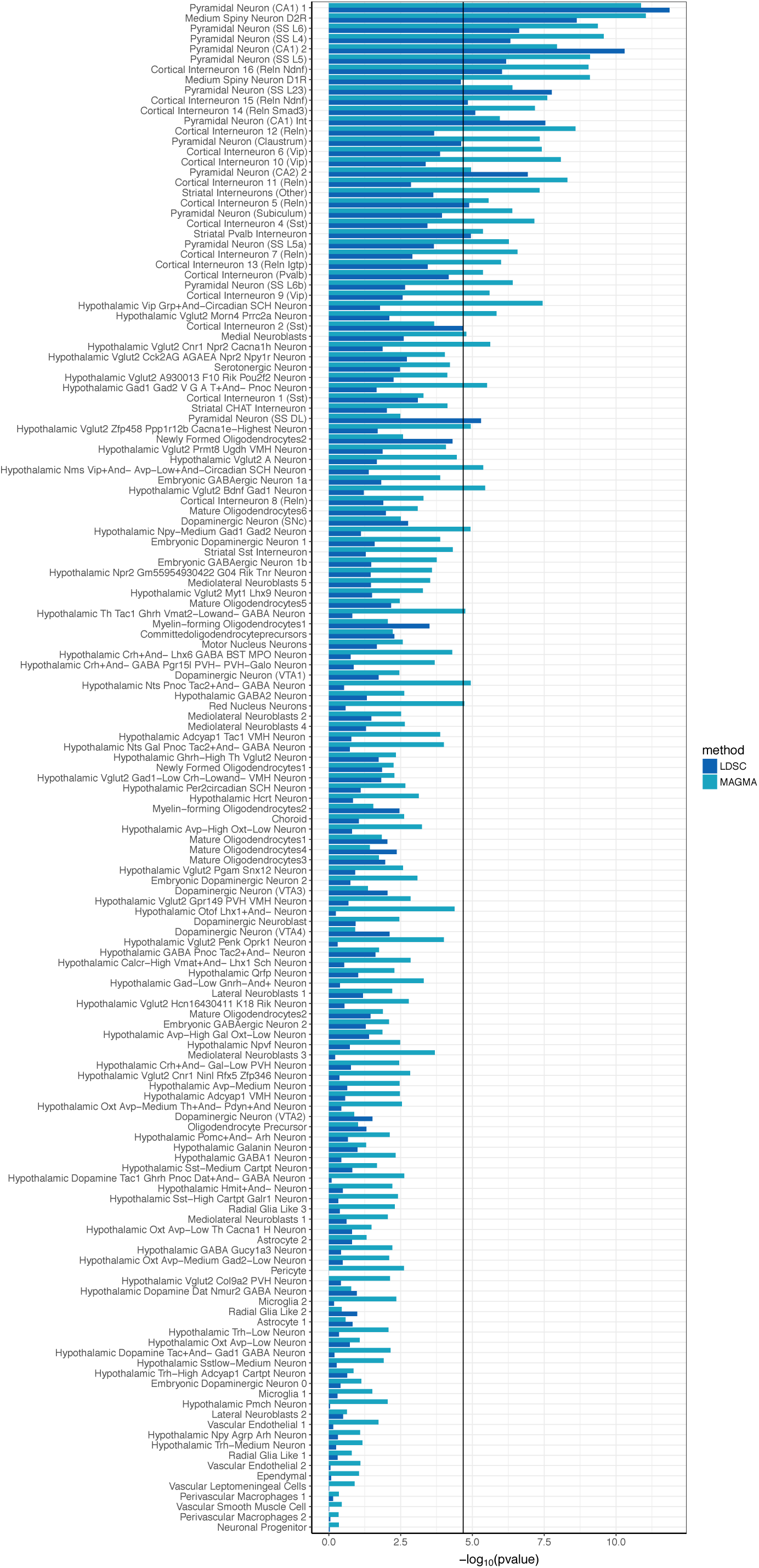
Association between brain cell types (KI level 2) and schizophrenia. Cell types are ranked by the minimum average rank of LDSC and MAGMA (minimum average P-values for cell types with equal average rank). The black line represents the Bonferroni significance threshold.

**Figure S7.**
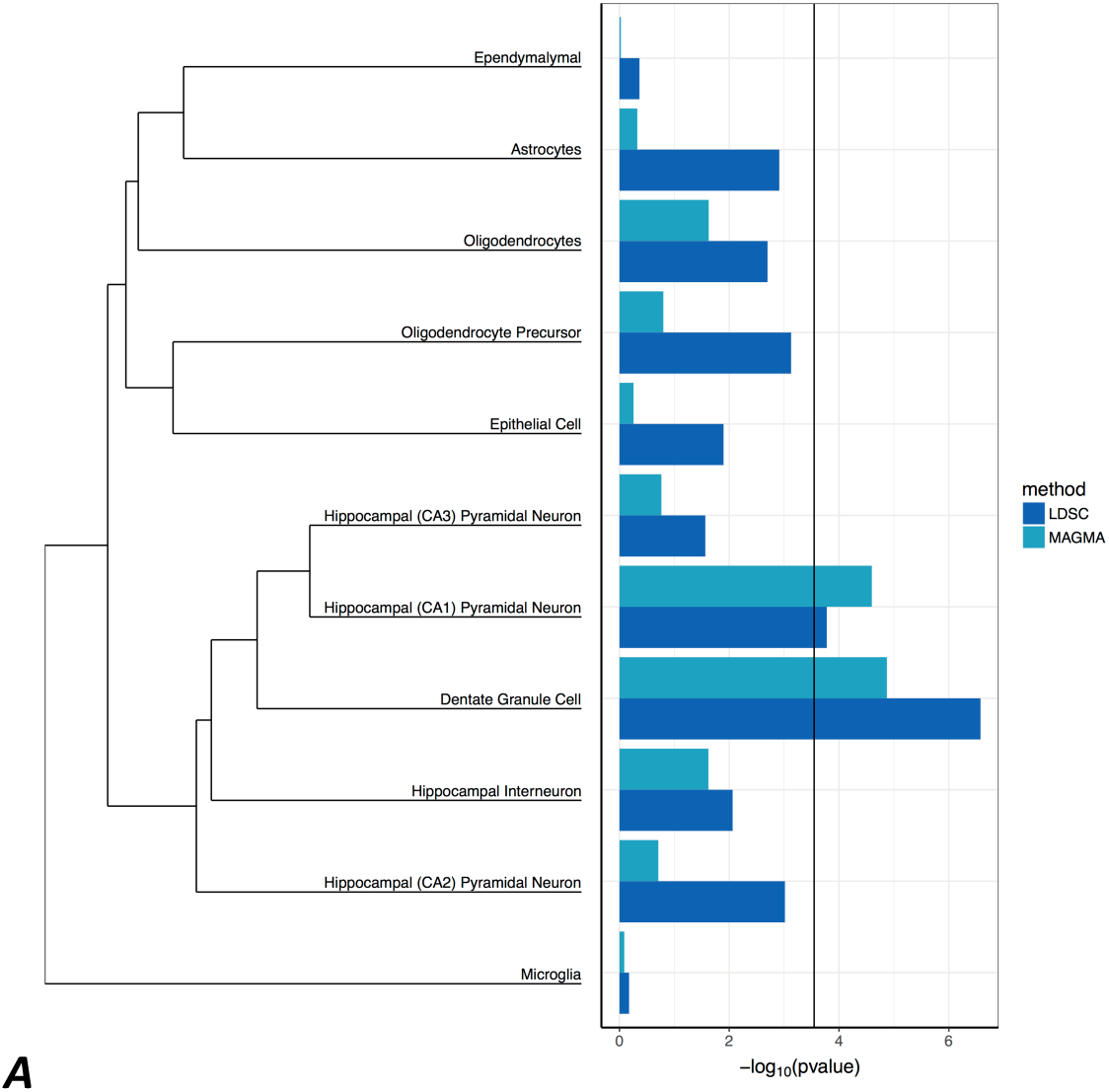
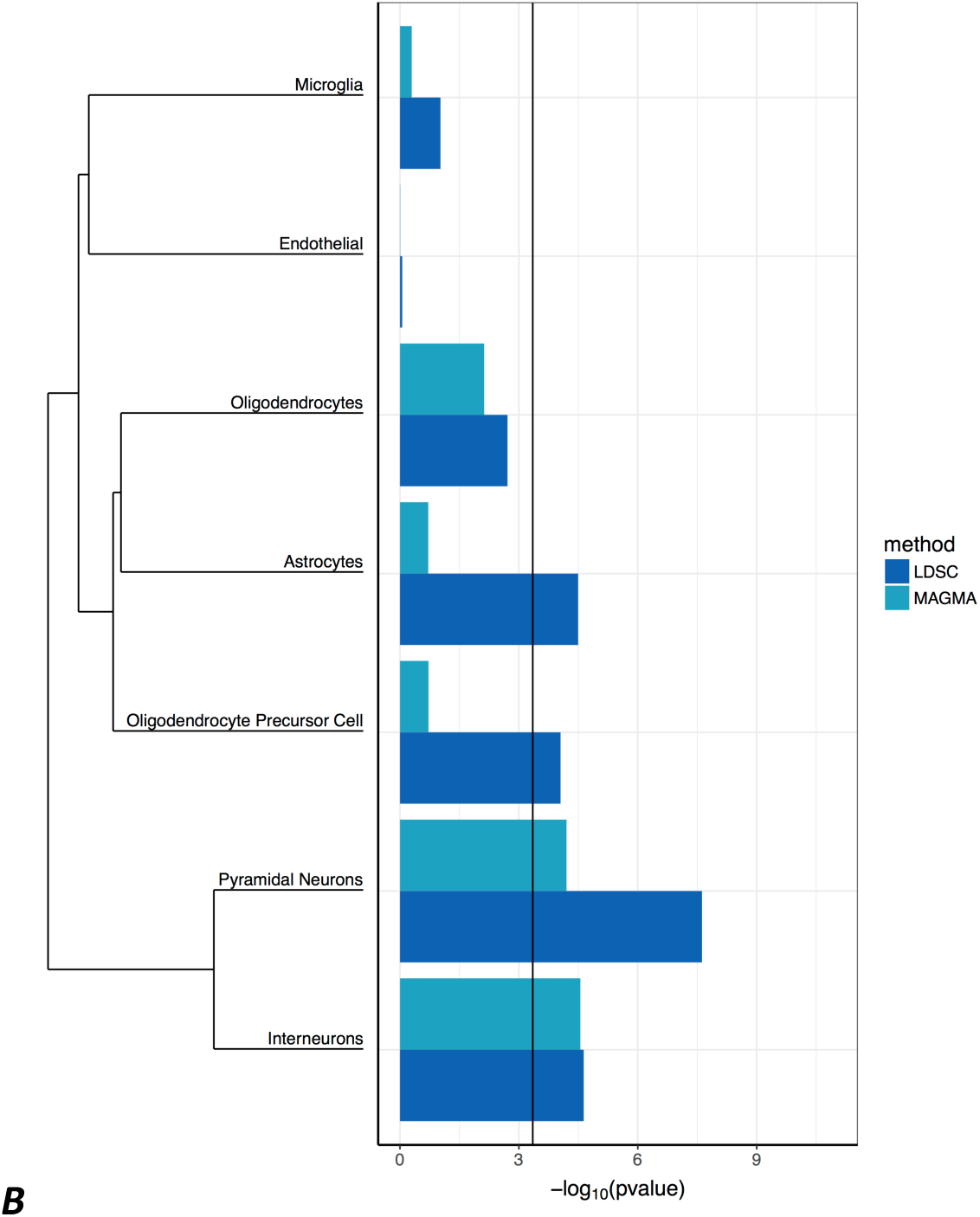
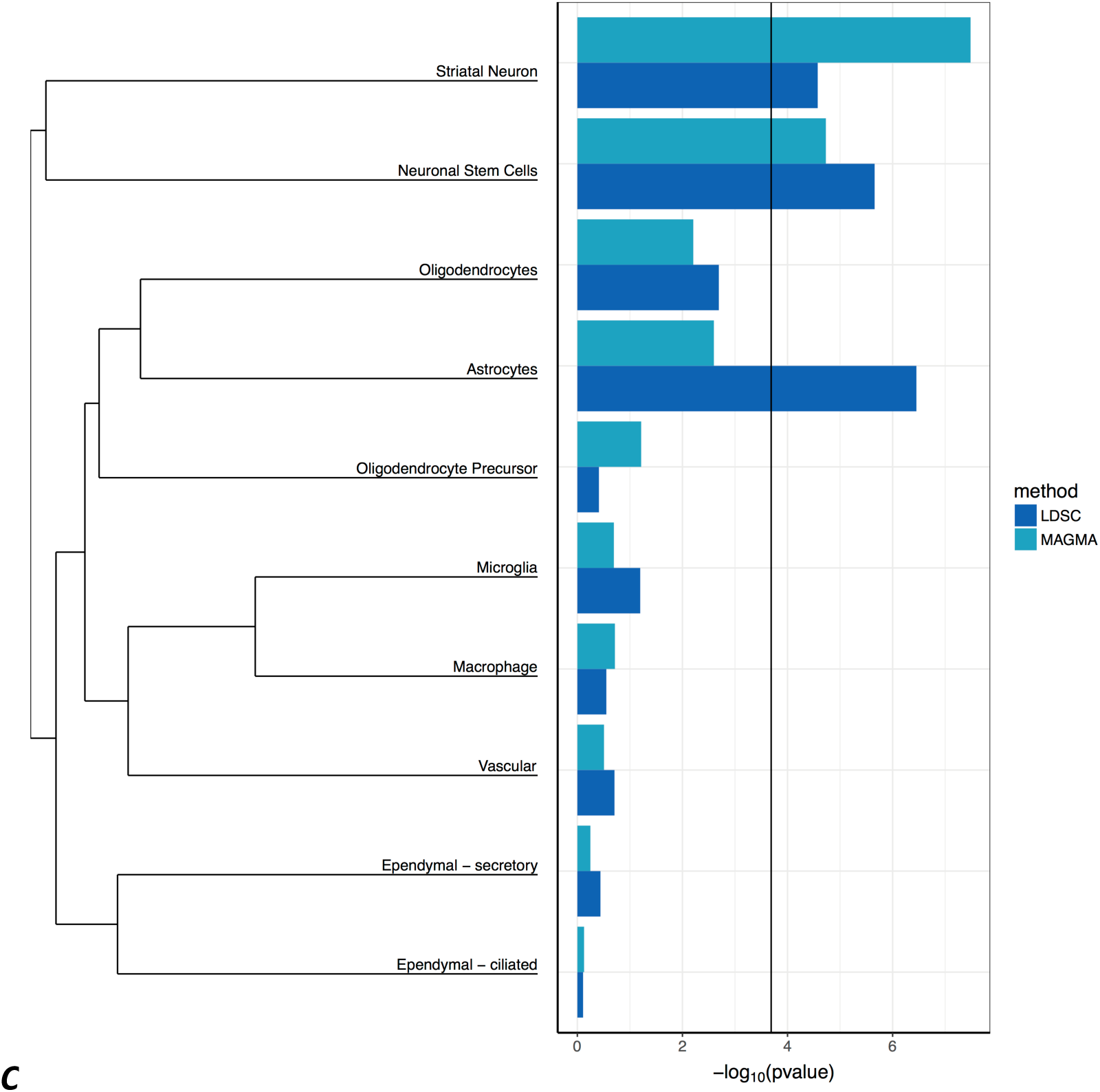
Confirmation of enrichment of common variant CLOZUK schizophrenia GWA results from the KI mouse data in independent mouse brain studies. The KI Level 1 findings connect the CLOZUK schizophrenia results to hippocampal CA1 pyramidal neurons, cortical pyramidal neurons, cortical interneurons, and medium spiny neurons. (A) snRNAseq from mouse hippocampus ^20^ showing enrichment of hippocampal CA1 pyramidal cells. (B) scRNAseq from mouse cortex ^21^ demonstrating enrichment of cortical pyramidal neurons and cortical interneurons. (C) scRNAseq from mouse striatum with enrichment of “striatal neurons”, which were predominantly medium spiny neurons ^19^. There was also enrichment for hippocampal dentate granule cells in Figure 6a and migratory neural precursors in striatum in Figure 6c, but these were not included in the larger KI dataset.

**Figure S8.**
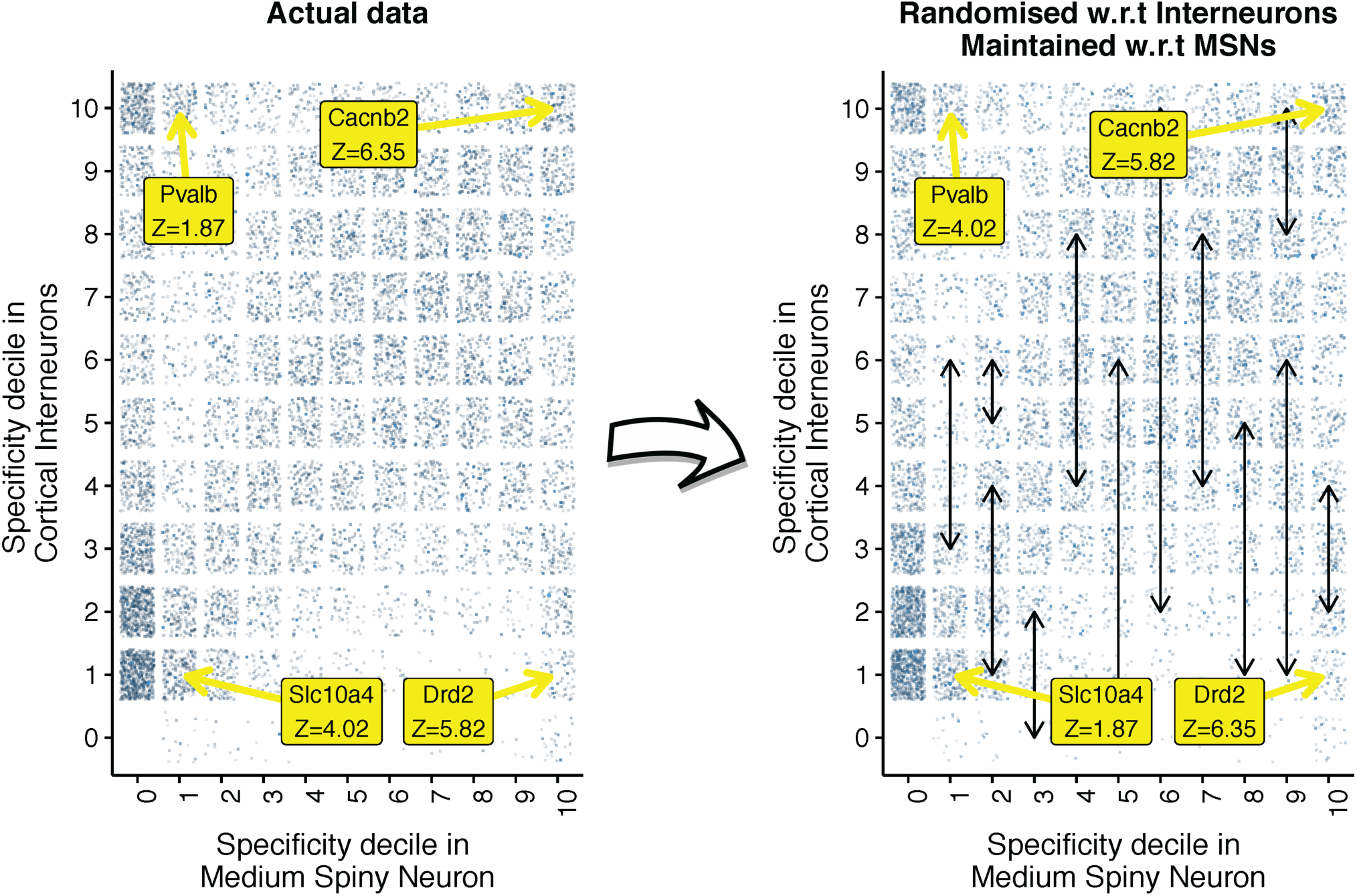
Schematic of the resampling approached used for conditional cell type enrichments shown in Figure 4a. The purpose of the resampling method was to enable testing of whether the schizophrenia enrichment detected for one cell type (here represented by cortical interneurons, INTs) is just an unavoidable result of a significant enrichment in a second cell type (here represented by Medium Spiny Neurons, MSNs). The plot on the left shows the correlation in specificity values between INTs and MSNs. Each point represents a single gene. Darker points have higher Schizophrenia CLOZUK association z-scores (calculated using MAGMA). The right hand plot shows how the z-scores are resampled within each MSN specificity decile, such that while the specificity values of each gene in MSNs and INTs remains the same, and the distribution of z-scores remains constant relative to each MSN decile, the distribution of z-scores is however randomized relative to INTs. Yellow boxes and arrows mark the location and z-scores of four genes: in the left hand plot, the z-scores are those found in the MAGMA genes.out file, while in the right hand plot the z-scores shown are resampled (hence the Pvalb scores has been switched with those of Slc10a4, as both are in the same specificity decile for Medium Spiny Neurons). For the analysis shown in Fig4a, this resampling of z-scores is performed 10000 times over and the relative enrichment of INTs in the left plot compared to that in the resampled right hand plot.

**Figure S9.**
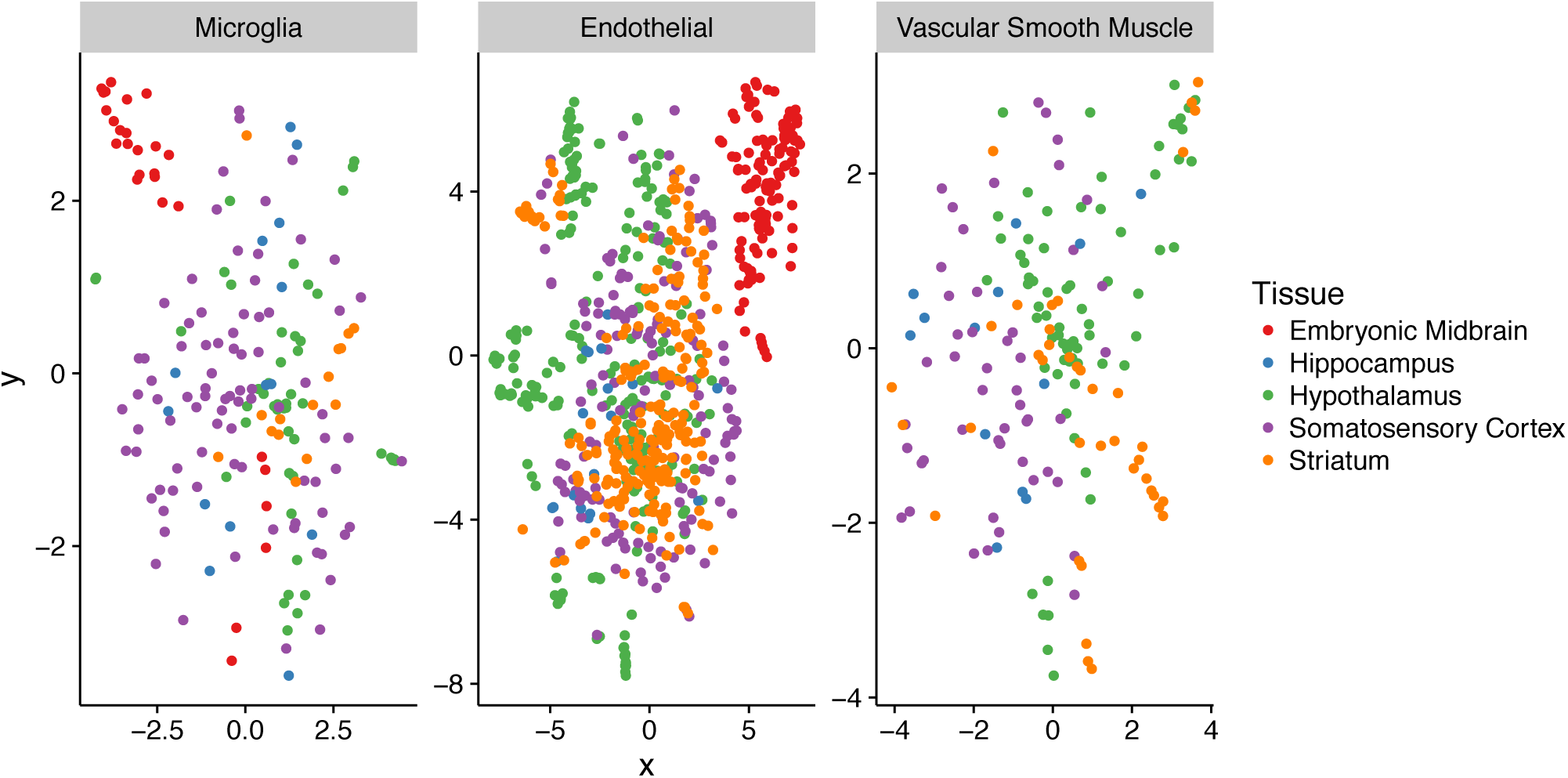
tSNE plot of three cell types shared between brain regions from the KI superset shows that they cluster together across regions. Microglia, Endothelial cells, and Vascular Smooth Muscle cells were selected on the basis that there is little prior expectation for them to have regional differences in expression. We found that embryonic midbrain cells cluster separately as is expected as they were obtained from embryonic tissue whereas all other samples were from adolescent mice. The cells from the other datasets were largely overlapping confirming that little to no batch effects exist in the data.

